# The marked diversity of unique cortical enhancers enables neuron-specific tools by Enhancer-Driven Gene Expression

**DOI:** 10.1101/276394

**Authors:** Stefan Blankvoort, Menno P. Witter, James Noonan, Justin Cotney, Cliff Kentros

## Abstract

Understanding neural circuit function requires individually addressing their component parts: specific neuronal cell types. However, not only do the precise genetic mechanisms specifying neuronal cell types remain obscure, access to these neuronal cell types by transgenic techniques also remains elusive. While most genes are expressed in the brain, the vast majority are expressed in many different kinds of neurons, suggesting that promoters alone are not sufficiently specific to distinguish cell types. However, there are orders of magnitude more distal genetic cis-regulatory elements controlling transcription (i.e. enhancers), so we screened for enhancer activity in microdissected samples of mouse cortical subregions. This identified thousands of novel putative enhancers, many unique to particular cortical subregions. Pronuclear injection of expression constructs containing such region-specific enhancers resulted in transgenic lines driving expression in distinct sets of cells specifically in the targeted cortical subregions, even though the parent gene’s promoter was relatively nonspecific. These data showcase the promise of utilizing the genetic mechanisms underlying the specification of diverse neuronal cell types for the development of genetic tools potentially capable of targeting any neuronal circuit of interest, an approach we call Enhancer-Driven Gene Expression (EDGE).

**Highlights:**

- Enhancer ChIP-seq of cortical subregions reveals 59372 putative enhancers.
- 3740 of these are specific to particular cortical subregions.
- This reflects the remarkable anatomical diversity of the adult cortex.
- Unique enhancers provide a means to make targeted cell-type specific genetic tools.

## INTRODUCTION

The mammalian brain is arguably the most complex biological structure known, composed of around 10^11^neurons in humans[1]. While this number is generally accepted, the same is not true for how many different *kinds* of neurons exist. Indeed, there is not even a clear consensus as to how to define a neuronal cell type: by morphology, connectivity, gene expression, receptive field type, or some combination of the above? If one takes the expansive view (i.e. all of the above), the numbers quickly become astronomical. For example, current estimates of retinal cell types range between 100 and 150[2], and dozens of cell types have been proposed for a single hypothalamic area based solely on which genes are expressed[3]. However, gene expression alone is a poor basis for defining cell types, because although most genes are expressed in the adult brain, the vast majority of them are expressed in many different cell types[4]. Identification of neuronal cell types is much more than an issue of taxonomy, it is crucial to understanding brain function. The past two decades have seen the development of revolutionary molecular tools which allow one to determine the precise connectivity of neurons[5, 6] as well as manipulate[7-9] and observe[10] their activity. Yet, the utility of these powerful tools is often limited by the inability to deliver them at the level of particular neuronal cell types. Almost all existing neuron-specific lines are either made by non-homologous recombination of minimal promoter constructs[5, 11, 12] or knocking the transgene into the native transcript[13, 14], made much easier by the advent of CRISPR-Cas. Both of these techniques depend upon the specificity of a native promoter, which can recapitulate the expression of the native gene, but that gene will almost always be expressed in multiple neuronal cell types[15-17].

Still, there must be *some* genetic basis for the remarkable diversity of neuronal cell types. Investigations of eukaryotic transcriptional regulation have revealed that spatiotemporally precise gene expression is achieved by the modular and combinatorial action of a variety of trans–acting factors (i.e. DNA-binding proteins) interacting with cis-regulatory elements (i.e. regions of noncoding DNA termed enhancers)[18]. While the exact number of enhancers remains unknown, estimates run into the millions[19, 20]. This is many times the number of genes or promoters, suggesting that the same gene is expressed in distinct cell types via the activation of different sets of enhancers. Enhancers may therefore enable the generation of molecular genetic tools that are more specific than what is possible using promoter-based methods. Indeed, many of the most specific neuronal driver lines are likely the result of random integration next to a highly specific enhancer[12, 21]. Fortunately, investigators studying the mechanisms of transcription have developed a variety of techniques enabling the identification of the enhancers active in any tissue sample[22-26]. We reasoned that because different cell types are found in different brain regions, enhancers active only in particular brain regions could enable the generation of region-and/or cell type-specific molecular genetic tools, an approach that we call Enhancer Driven Gene Expression or EDGE (Figure 1).

**Figure 1.**
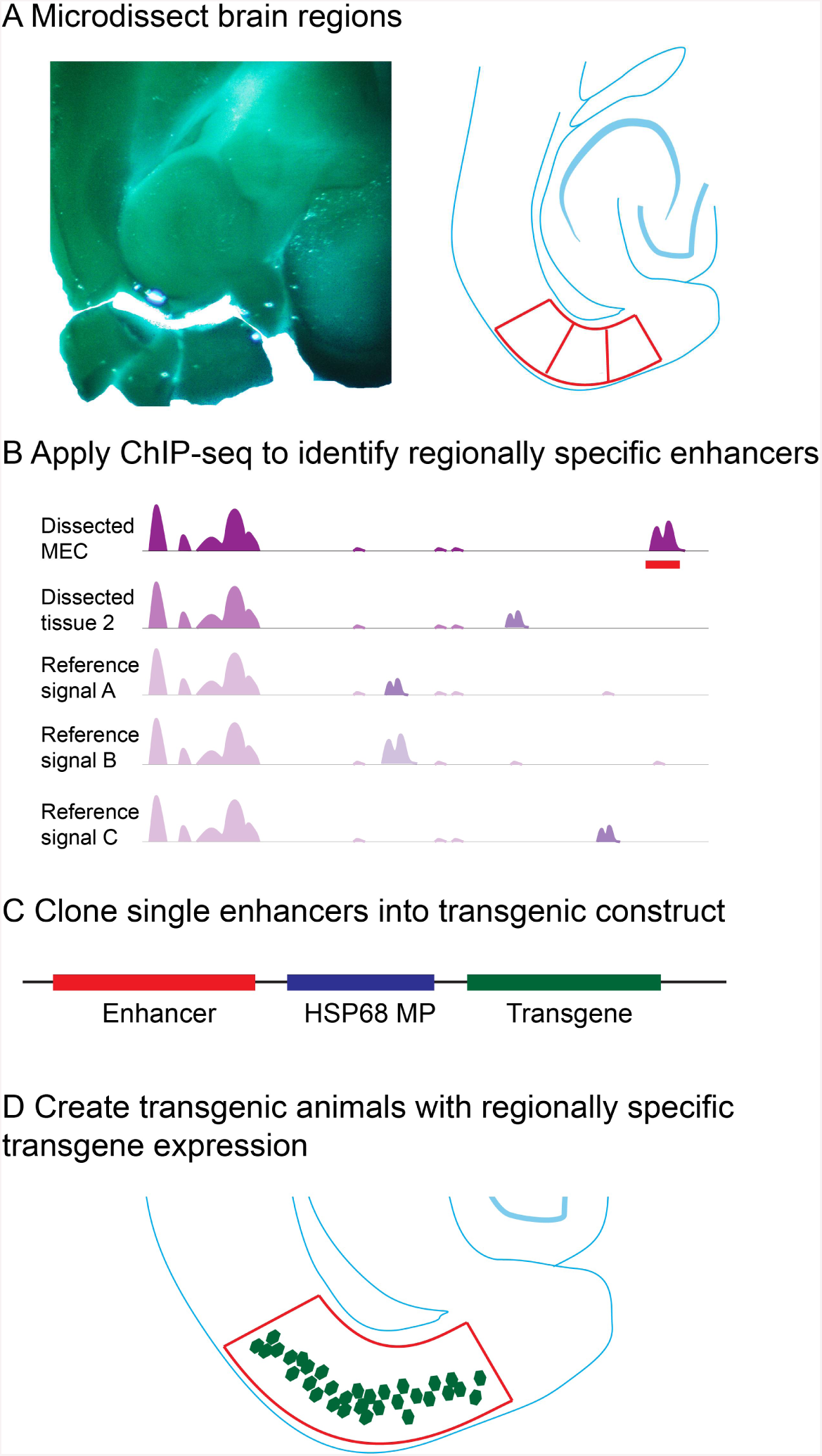
Experimental summary of Enhancer Driven Gene Expression (EDGE). (A) Samples of brain regions of interest are microdissected by hand. (B) ChIP-seq is performed on these samples and genome-wide H3K27ac and H3K4me2 signals for each sample are compared to reference signals and signals from the other samples. Bioinformatic analysis algorithms output unique peaks as potential region-specific enhancers (red bar). (C) Single putative enhancers are cloned into constructs containing a heterologous minimal promoter to drive transgene expression. (D) Following pronuclear injection of these constructs, the resulting founder mice are crossed to reporter lines and evaluated for desired expression patterns. See also Figure S1.

## RESULTS

### Enhancer ChIP-seq of cortical subregions reveals a striking diversity of unique enhancers

Because promoter based techniques generally lead to gene expression throughout the telencephalon, we specifically targeted closely related subregions of cortex in the hopes of obtaining regionally specific tools. The following brain regions from two adult (P56) male C57BL6J mice were microdissected (for details see methods and Figure S1): the medial entorhinal cortex (MEC), the lateral entorhinal cortex (LEC), the retrosplenial cortex (RSC), and the anterior cingulate cortex (ACC). Each mouse was processed separately and the samples were used as biological replicates for further analysis. We performed ChIP-seq on homogenized tissue against the active-enhancer-associated histone modifications H3K27ac and H3K4me2 for samples of each of the four brain regions. The regions enriched for H3K27ac reproducibly identified similar numbers of active promoters and distal cis-regulatory sequences between two replicates of each brain subregion (Figure 2A). Nearly 90% of all active promoters were identified in at least two subregions with the remainder being active in only one subregion (17032 total, 2045 unique).

**Figure 2.**
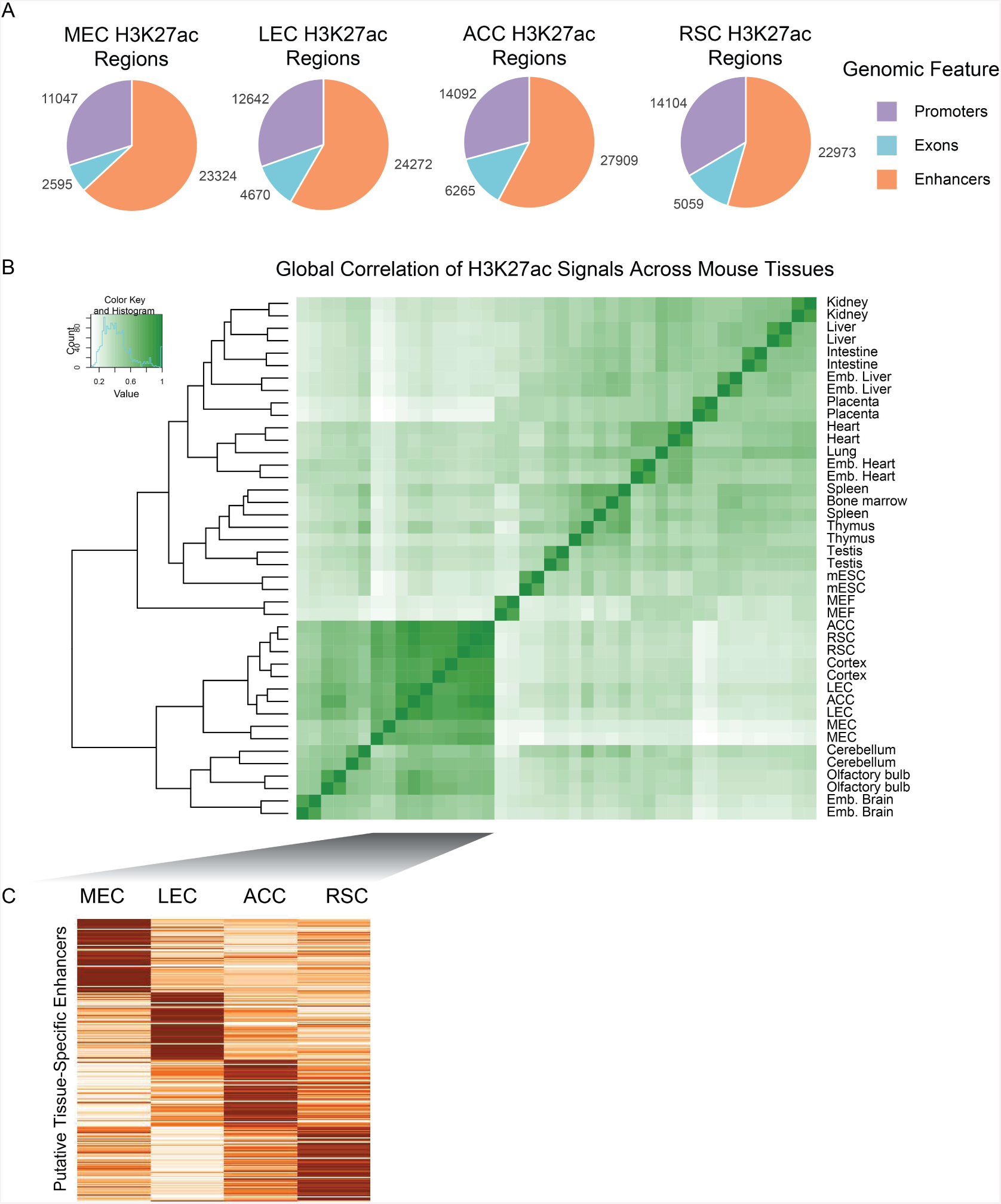
ChIP-seq reveals a striking diversity of unique and novel enhancers in different cortical subregions. (A) Pie charts showing the proportions (and numbers) of distinct active genomic elements identified by H2K27ac ChIP-seq of the 4 cortical subregions. These numbers are roughly similar to those found by ChIP-seq of other organs. (B) Dendrogram (left) and correlation matrix of the H3K27ac signals (right) from replicates of the cortical subregions dissected in this experiment versus those from ENCODE were used for subtraction. Note the relatively high correlation of replicates (except ACC) and clustering of signal from cortical tissues. (C) Heatmaps showing some of the tissue-specific putative enhancers identified in the microdissected cortical subregions. See also Figure S2 and Data S1.

When we analyzed more distal sites (>5kb from a transcriptional start site) we identified a total of 59372 reproducibly active enhancers in at least one subregion. Of these 31% were only identified in a single cortical subregion (18185 unique relative to other subregions). Surprisingly the number of subregion specific enhancers in the cortex was similar to the number of total enhancers active in any single tissue in the body thus far investigated[20, 27]. Furthermore, 81% (48077) of enhancers identified in these subregions were not identified in bulk cortex tissue, presumably due to signal-to-noise ratios. The fact that so many novel and unique enhancers were isolated from a tiny minority of cortical regions demonstrates the potentially vast repertoire of enhancers active in the brain.

Interestingly, when comparing the total number of reproducible peak calls in these 4 cortical subregions (59372) to the number identified in bulk cortex treated in the same way (13472), the number of putative active enhancers one obtains from the four cortical subregions is far greater than what one obtains from the entire cortex, even though these four cortical regions compose only a small minority of the entire cortex. Of course, this is comparing 4 pooled samples to a single sample, but each of the individual samples gives numbers similar to bulk cortex (Figure 2). In our view, the most likely explanation for this superficially puzzling result is a reduction in signal to noise ratio when pooling heterogeneous sets of tissues for ChIP-seq. This would tend to favor those enhancers that are expressed throughout many cortical subregions at the expense of more specific ones. In support of this, 89% of cortical enhancers were found in one or more cortical subregions, and 78% were found in at least 2 cortical subregions. Compare this to the fact that fully 31% of the enhancers we found in our subregions were specific to that single subregion.

While many of these enhancers identified by peak calls alone are specific to this small number of cortical subregions the goal of this study was to identify very specific regulatory sequences with limited activity within other regions of the brain as well as the rest of the body. To ensure the identification of such sequences and exclude regions with weak activity elsewhere we expanded our comparisons to include a variety of published mouse adult tissues and cultured cell types[27]. We first identified active putative enhancers in these additional mouse samples and merged them to create a unified set of enhancers for consistent comparisons across all samples. We then extracted normalized H3K27ac counts at 108299 discrete regions from the subregions profiled in this study as well as those from 17 mouse ENCODE samples[27]. Hierarchical clustering of samples revealed two main groups of mouse tissues: neuronal and non-neuronal (Figure 2B). Amongst non-neuronal tissues, the strongest correlations were observed in developmental stages of heart and tissues that make up the immune system: bone marrow, thymus, and spleen. In neuronal tissues the four cortical subregions profiled here were well correlated across all enhancers assayed but clustered distinctly from cerebellum, olfactory bulb, and embryonic brain.

We then utilized k-means clustering to identify enhancers that were significantly more active in each cortical region versus each other (Figure 2C) and the other 17 mouse tissues. Those enhancers that were identified as most specifically active in a given cortical subregion were then further filtered to ensure that they were never identified by peak calling in any other mouse tissue. Even though this does not exclude identification of enhancers as unique while they are actually highly enriched, it does increase the specificity and thus the chances of identifying unique enhancers. This stringent analysis yielded 165 to 1824 novel and unique putative distal enhancers for each cortical subregion (Figure 2C, Data S1). We then assigned these novel enhancers to putative target genes based upon the GREAT algorithm[28]. Gene ontology analysis suggests these novel enhancers are enriched near genes associated with a variety of neuronal functions (Figure S2). We prioritized these novel putative enhancers based on specificity of the H3K27ac signal relative to other regions and conservation across 30 species. We then cloned a subset of them specific to the entorhinal cortices (EC) upstream of a heterologous minimal promoter driving the tetracycline transactivator (tTA[29]) for transgenesis (Figure S3).

### Region-specific enhancers drive transgene expression in targeted cortical subregions

Of course, just because a sequence is identified by ChIP-seq does not mean that it is a valid enhancer, let alone that it can drive region-or cell type-specific transgene expression. Even a single case of expression in a particular tissue type is not sufficient because one can obtain specific transgene expression by randomly inserting a minimal promoter/reporter construct into the genome. This technique is known as an “enhancer trap” because it relies upon random insertion near a native enhancer to drive the transgene expression[12, 30]. To ensure that the expression pattern comes from the enhancer construct and not from the insertion site, the standard way to validate a putative enhancer is to show that at least three distinct transgenic embryos (with three distinct random insertion sites) have similar expression patterns[26]. We therefore injected enough oocytes to get at least three genotypically-positive founders for each putative enhancer construct. But, since our aim was to generate modular genetic tools rather than simply to validate the enhancers, we could not sacrifice the founders to validate the enhancer as is typically done. Instead, the founders were crossed to tTA dependent reporter mice for visualization of expression patterns.

We selected 8 (notionally) MEC-specific and 2 LEC-specific enhancers for transgenesis. Transgenesis via pronuclear injection is not an efficient process because it involves random integration into the genome. While one typically only publishes the ones that work, it is worth specifying what issues have to do with transgenesis in general versus using EDGE. When making any transgenic line, some founders do not successfully transmit the transgene to offspring, while others fail to express presumably due to negative insertional (a.k.a. positional) effects. For these reasons, only 45 lines derived from 105 genotypically-positive founders expressed in the brain when mated to a tetO reporter line, a number that is typical regardless of the injection construct. Notably, nearly all of them (41) expressed the reporter in the EC, including at least one from each of the 10 enhancer constructs (Figure S4 and Table S1). Since an enhancer trap would lead to random expression patterns, this alone suggests that the specificity of expression comes from the transgenic enhancer. At least as compelling is the fact that when we obtained multiple distinct founders with a given enhancer construct, almost all of them had similar expression patterns (see Figure S5 for examples).

Figure 3A shows an example of the results of our bioinformatic analysis for one of the eight MEC enhancers (MEC-13-81, see methods for nomenclature) which GREAT associated with the gene *Kitl*. Note that the promoter region (vertical yellow band) is a strong peak in all brain regions, consistent with expression of the *Kitl* mRNA throughout the brain (Figure 3B). The same is true for other putative enhancers (horizontal black bars). In contrast, the downstream enhancer peak used for transgenesis (MEC-13-81, Figure 3A blow-up), while not as strong as some of the other peak calls, is greatly enriched in MEC. Figure 3C shows the result of crossing the transgenic line MEC-13-81B to an tetO-ArChT payload line[31]. Remarkably, even though the *Kitl* promoter expresses throughout the brain (including multiple layers of the EC, Figure 3B), the tetO-ArChT payload is confined to layer II of MEC (Figure 3C). In other words, one can obtain highly specific targeted gene expression from regionally specific cis-elements of non-specific genes.

**Figure 3.**
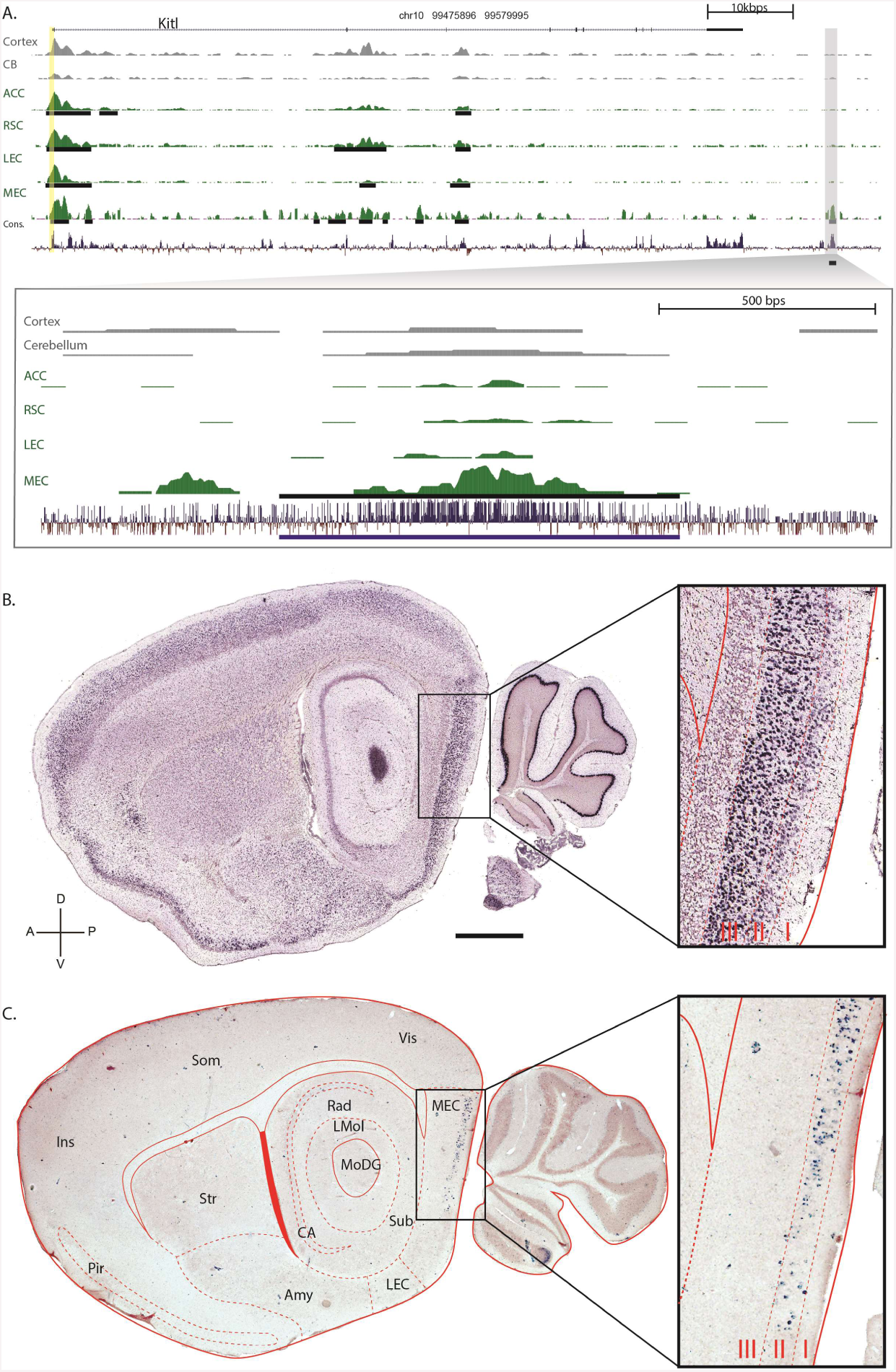
The enhancers of non-specific genes drive region-specific transgene expression. (A) A genomic view of one of the 165 MEC-specific enhancers yielded by ChIP-seq analysis. The top panel indicates the location and coding regions of Kitl as well as H3K27Ac signal for two regions from Roadmap epigenome (Cortex and Cerebellum), the four regions we analyzed (ACC, RSC, LEC and MEC), and conservation over 30 species. The vertical yellow column indicates the promoter region upstream of the transcriptional start site. Peak calls, are denoted by the black horizontal lines. The specific genomic region containing the enhancer (MEC-13-81) is blown up in the bottom panel. (B) ISH (brain-map.org) of Kitl, the gene associated with enhancer MEC-13-81 shows expression throughout cortex, hippocampus and cerebellum. (C) tTA dependent transgene Arch driven by the enhancer (ranked number 81) is expressed in MEC LII. Scalebar is 1000µm. Sagittal plane, Dorsal-Ventral and Anterior-Posterior axis are indicated. Abbreviations are: Ins: insular cortex, Som: somatosensory cortex, Vis: visual cortex, Pir: Piriform cortex, Str: striatum, Amy: amygdala and associated regions, Rad: stratum radiatum of the hippocamlus, LMol: molecular layer of the hippocampus, CA: both cornu ammonis fields of the hippocampus, sub: subiculum, MoDG: moclecular layer of the Dentate Gyrus, LEC: lateral entorhinal cortex, MEC: medial entorhinal cortex, Layers I, II and III of the MEC are indicated in the blow-up to the right. See also Figure S3, S4, S5, S6 and Table S1.

The same basic result of highly specific expression from single enhancers of non-specific genes was also true for 4/8 MEC-and 2/2 LEC-specific enhancer constructs we injected, although the correspondence between the ChIP-seq signal and the expression was not always as tight. Figure 4 compares the expression patterns of representative transgenic driver lines made with other injection constructs containing either MEC-specific enhancers (Figure 4A to 4C, right column) or LEC-specific enhancers (Figure 4D and E, right panel. Extended medial-lateral of sections range in Figure S4A-F) compared to the expression pattern of the presumed associated native gene (Figure 4 left column). Note that while each associated gene is broadly expressed in the brain, the transgenic lines all express more or less specifically in the brain region the enhancers were isolated from. When crossed with broadly expressing tTA lines (such as CaMKIIa [11]), these tetO payload lines express broadly (see references for published lines and Figure S6 for our as-of-yet unpublished tetO-GCaMP6 line). These data show that one can obtain targeted region-specific (and possibly even cell type-specific) expression from elements of a non-specific promoter by using one of its region-specific enhancer to drive a heterologous core promoter. Even those enhancers that were less specific still gave rise to lines that were enriched in the EC relative to the expression of the native gene (Figure S4G-J). This in effect solves the problem that most genes are expressed in multiple cell types in the brain: using EDGE one can dissect out the individual genetic components which underlie the expression of a “nonspecific” gene in multiple cell types.

**Figure 4.**
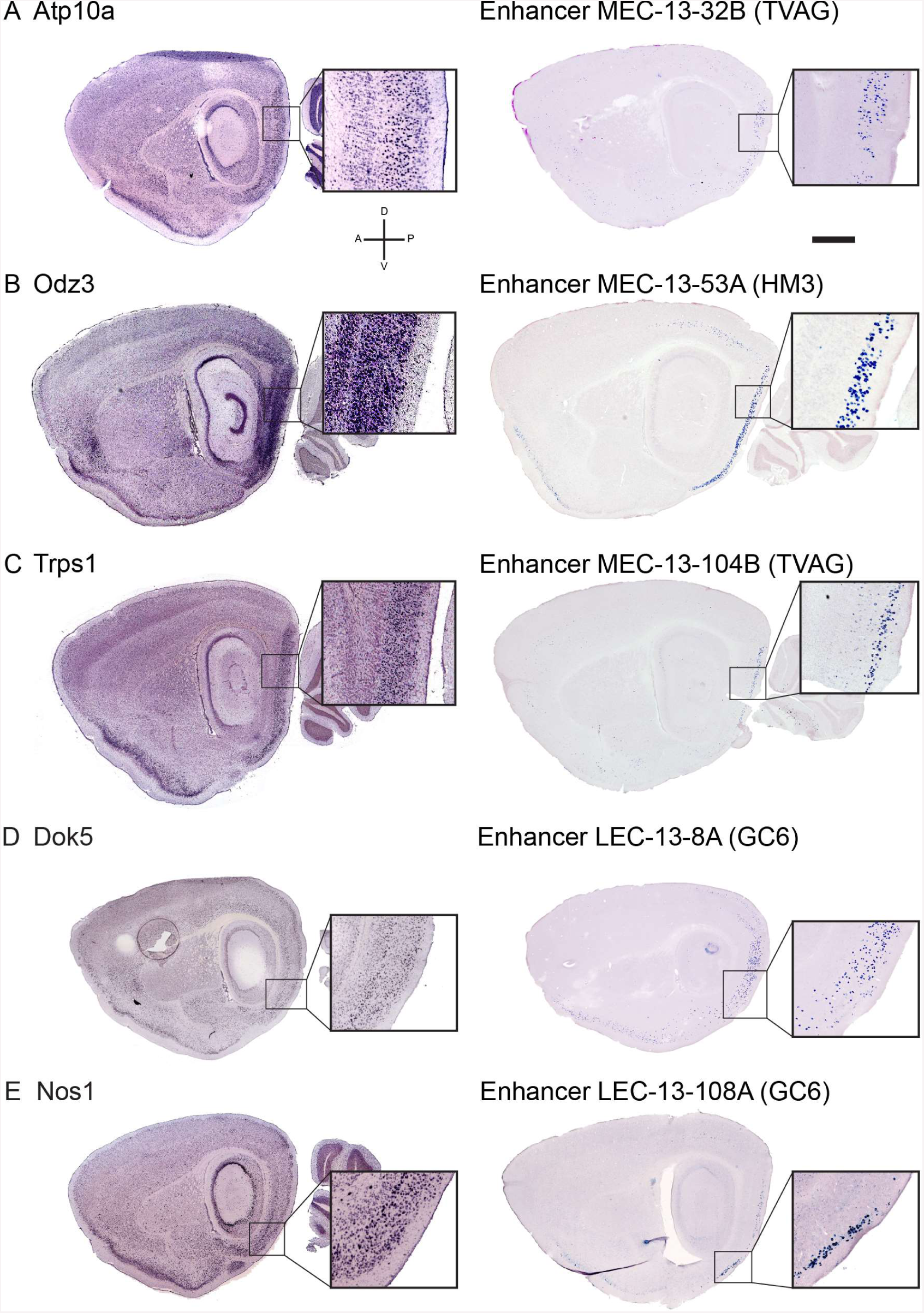
Distinct MEC-specific enhancers drive transgene expression in distinct sets of cells in MEC. (A through E, left column) ISH showing expression patterns of native genes associated with EC-specific enhancers. (A through E, right column) ISH showing EC-specific expression of transgenes driven by the corresponding EC-specific enhancers, tTA driven transgenes in parentheses. ISH for the native genes from brain-map.org. Scalebar in A is 1000µm. See also Figures S4, S5 and S6 and Table S1.

### Region versus cell type-specific expression?

The above results show that subregion specific expression can result from subregion specific enhancers. Whether such enhancers drive expression in specific cell types in the targeted brain region is a more difficult question to answer, in large part because there is no consensus as to the number of cell types in the brain or how to classify them. However, there are indications that some these enhancers can specify particular cell types, at least to the level of granularity current knowledge permits. First, the different EC enhancers tend to drive expression in different layers of the EC (Figure 3 and 4 and S7), and neurons in different cortical layers are almost by definition different cell types. By the same logic, some of these enhancers are clearly not cell type-specific (Figure S4G-J). Since four of the enhancers drive expression in layer II, this raises the question of whether they specify the same cell type, or distinct biological subpopulations. We therefore investigated the expression of immunohistochemical markers used to characterize cell types of EC in two layer II expressing lines derived from MEC-specific enhancers (Figures 5A, B and 6A, B). The underlying logic is that if the two distinct enhancers drive transgene expression in subsets of the exact same cell type(s), they should both express the same proportions of neurochemical markers. Neither of the two enhancers appear to drive expression in inhibitory neurons (Figure 5I-L and 6I-L), so the question becomes whether they express in different types of excitatory neurons. Excitatory neurons in EC layer II are typically further subdivided into reelin positive cells and calbindin positive cells[32]. Line MEC-13-53A expressed exclusively in reelin+ neurons (Figure 5C-H, L), while line MEC-13-104B roughly corresponds to the relative densities of the two celltypes (Figure 6C-H, L). Thus it appears that MEC-13-53A is a stellate cell specific enhancer, whereas MEC-13-104B is found in both neurochemical kinds of excitatory cells of layer II described to date. This means that some enhancers specify different subsets of cells even within a single cortical layer, showing the potential of enhancers to distinguish between cell types with a finer granularity than possible with native promoters. Of course, the functional significance, if any, of these subsets of cells remains to be demonstrated.

**Figure 5.**
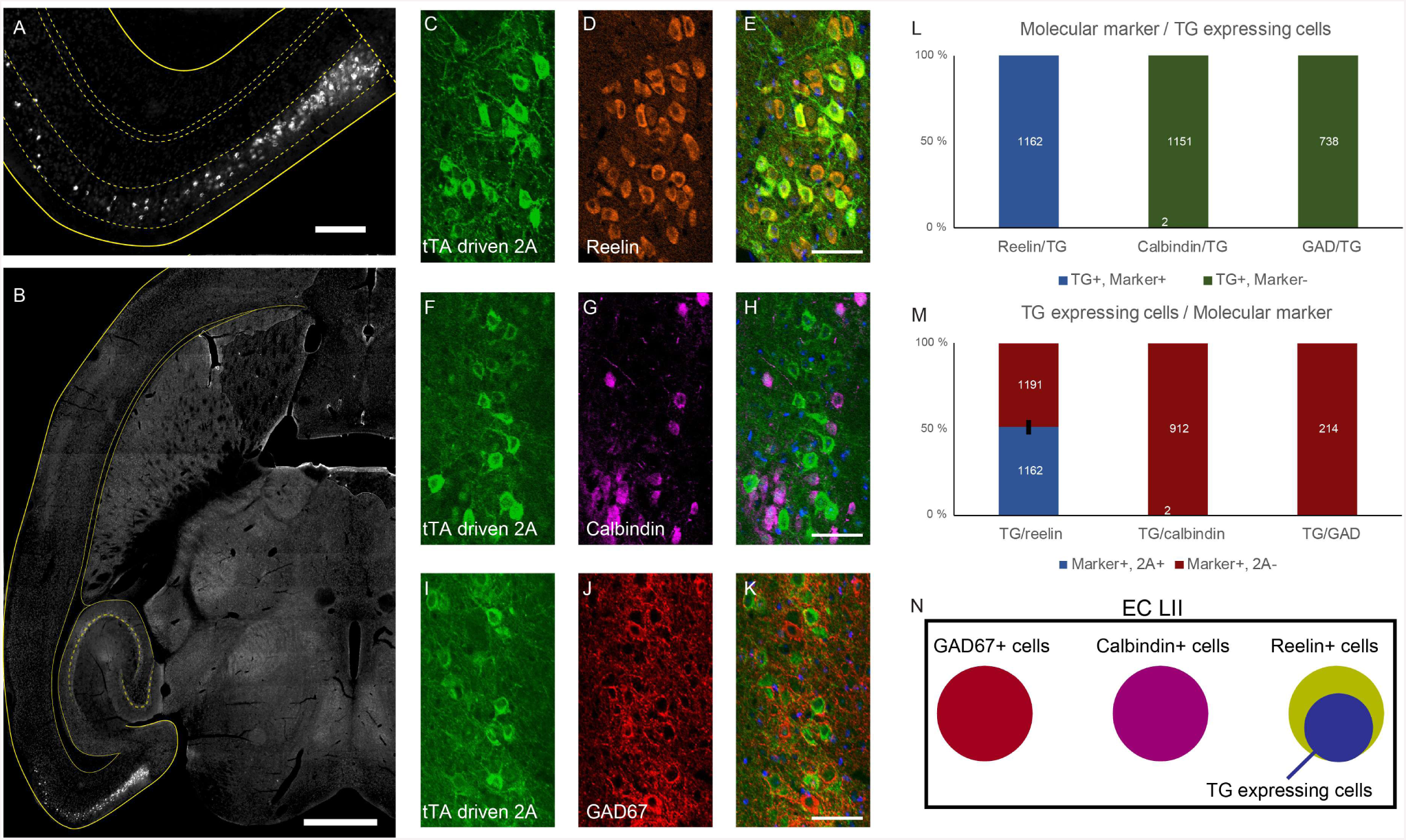
Single enhancers can drive expression in histochemically-defined subsets of MEC LII cells. (A,B) Horizontal section of a mouse cross between MEC-13-53A and TVAG. Immunohistochemical transgene detection with anti-2A Ab shows layer II EC-specific expression. (C,F,I) Anti-2A histochemistry; (D) Anti-Reelin; (G) Anti-Calbindin;(J) Anti-GAD67; (E,H,K) Overlays of the two signals, each row is the same section. (L) 100% (1162/1162 counted cells) of transgenic cells co-localize with Reelin but there is essentially 0% co-localization with calbindin (2/1151) and GAD67 (0/738). (M) 49.4% (1162/2353) of all Reelin positive cells were positive for the transgene, essentially none of the other cell populations had any transgene expressing cells. Total numbers of cells counted in white. (N) Schematic summary of the data in C to M. Scale bars are 1000µm in B, 200µm in A and 50µm in C-K. In all graphs bars show the mean ±SEM. See also Figure S7.

**Figure 6.**
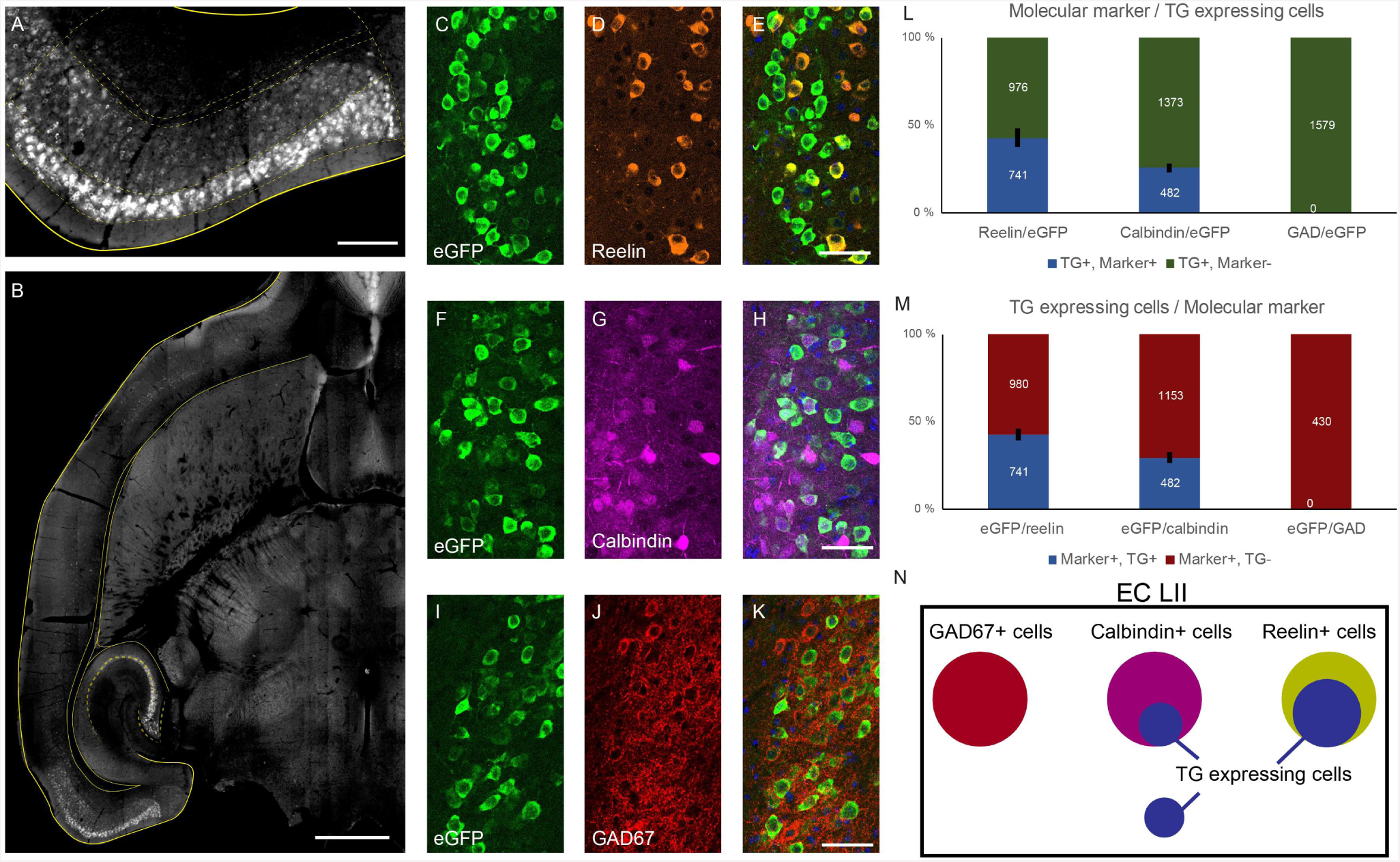
Different single enhancers can drive expression in histochemically-distinct subsets of MEC LII cells. (A,B) Horizontal section of a mouse cross between MEC-13-104B and tetO-eGFP. Immunohistochemical transgene detection with anti-GFP Ab shows expression in layer II of the EC. (C,F,I) Anti-GFP histochemistry; (D) Anti-Reelin; (G) Anti-Calbindin; (J) Anti-GAD67; (E,H,K) Overlays of the two signals, each row is the same section. (L) 43.1% (741/1717 counted cells) of transgenic cells in layer II of the EC co-localize with Reelin while 26% (482/1855) of them co-localize with calbindin. 0% (0/1579) co-localize with GAD67. (M) 43.1% (741/1721) of all Reelin positive cells in layer II of the EC were positive for the transgene and 28.5% (482/1635) of all Calbindin positive cells in layer II of the EC were positive for the transgene, while 0% (0/430) of the GAD67 positive population had any transgene expressing cells. Total numbers of cells counted in white. (N) Schematic summary of the data in C to M. Scale bars are 1000µm in B, 200µm in A and 50µm in C-K. In all graphs bars show the mean ±SEM. See also Figure S7.

**Figure 7.**
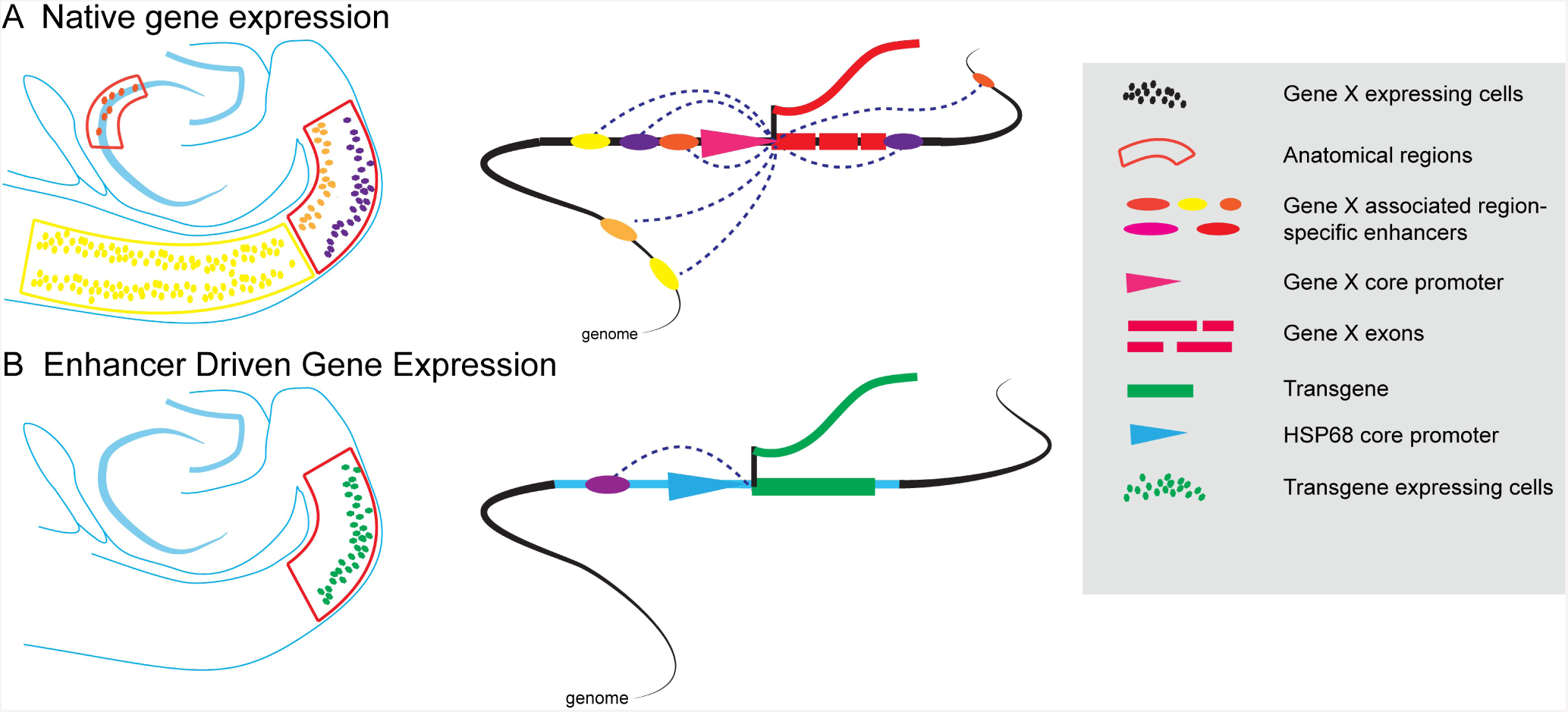
Schematic of putative genetic basis for EDGE technology. (A) Native gene expression: A gene “X” is expressed in multiple cell types in distinct brain areas. Expression in each cell type is driven by distinct sets of color-coded active enhancers acting upon the native core promoter (pink triangle). Promoter-based methods of transgene expression such as BAC transgenesis and Knock-ins respectively include several or all of the native enhancers, thereby recapitulating some or all of the expression pattern of the native gene. (B) Enhancer-Driven Gene Expression: a single active enhancer isolated from a particular brain region drives transgene expression from a heterologous minimal promoter (blue). This leads to transgene expression that is restricted to a particular region-specific subset of the cell types that the native promoter expresses in, greatly increasing the anatomical specificity relative to promoter-based methods or the native gene.

## DISCUSSION

We demonstrate the existence of thousands of previously undescribed putative enhancers uniquely active in targeted cortical subregions of the adult mouse brain. We took a small subset (10/3740) of the enhancers that were specific to the EC and combined them with a heterologous minimal promoter to make transgenic mice expressing the tTA transactivator. When crossed to tetO payload lines, we obtained transgene expression specific to the EC, and possibly even particular cell types in the targeted region. The genes that these enhancers (presumably) act upon are nowhere near that specific. Most genes express in multiple cell types in the brain. Since there are only around 24.000 genes (and around 46.000 promoters[19]), but estimated millions of putative enhancers[19, 20], this implies that the same gene is expressed in different cell types by using different sets of enhancers acting upon the same promoter. In turn, this suggests that there may be a genetic diversity in the brain beyond most estimates of the number of distinct neuronal cell types in the cortex[33-36]. Moreover, this provides a strategy to make genetic tools with far greater cell type and regional specificity of expression than promoter-based methods, by far the dominant means to generate neuron specific transgenic animals to date.

### EDGE is a method to create neuron-specific tools for targeted brain regions

While the above discussion illustrates the power of this technique, it is important to be clear about what is and is not novel about what has been presented here. A variety of forms of enhancer ChIP-seq have existed for roughly a decade[22, 26], and the general concept that the same gene is expressed in different tissues by the use of different enhancers is even older[30]. Hundreds of thousands of putative enhancers have already been identified in the mouse genome by dissection of distinct tissues (including cortex) followed by ChIP-seq[20, 27]. Indeed, a molecular geneticist in the transcription field may find the results presented here unsurprising, as generation of a transgenic animal is how putative enhancers are biologically verified, although the transgenic founders are typically killed in the process[22, 25, 26, 37]. In short, we have not invented any novel techniques, but we demonstrate how the application of these existing technologies to the adult brain could potentially provide systems neuroscientists with a means to make cell type-specific tools for any brain region of interest, greatly facilitating the study of the functional circuitry of the adult brain.

Putting our results into context requires discussing the rich literature that inspired our approach. A variety of recent papers used various techniques to suggest a highly diverse chromatin landscape in the adult brain, indicative of a diversity of enhancers. One group has performed ChIP-seq on 136 different dissected human brain regions, obtaining over 80.000 putative enhancers[38]. Another group has used ATAC-seq to profile open chromatin in transgenically-defined excitatory cells from different layers of the mouse visual cortex[39]. They found a diversity of putative cis-acting sequences even within single layers of a single type of cortex, implying distinct classes of cells. Finally, using single cell methylomes, Luo et al. have shown that neuron type classification is supported by the epigenomic state of regulatory sequences[40]. Nonetheless, in none of these cases were these putative enhancers biologically verified, nor used to make molecular genetic tools, which is the point of this paper.

Conversely, many enhancers derived from the developing brain have in fact been biologically verified, and even used to make transgenic lines and viruses[41]. Evolutionarily conserved single enhancers demonstrably label specific subsets of cells during development[25, 26, 37, 42], with different subsets active in different developmental epochs[43]. Of particular interest is a pair of papers from the Rubenstein lab examining the activity of enhancers derived from the developing (E11.5) telencephalon. They made CreER lines from the pallium (14 lines[44]) and subpallium (10 lines[45]) to illustrate the fatemaps of the telencephalic subdivisions by comparing expression patterns at several timepoints during development and young adulthood. By examining in vivo transcription factor occupancy they showed that broadly expressed transcription factors interact with far more specific enhancer elements[44].

Taken together, all these studies provide part of the basis for what is presented here. However, their focus was on the transcriptional and developmental mechanisms of neural cell fate relatively early in development. As these and other studies demonstrated, every neuroepithelial cell present at this time will have many daughter cells which will further differentiate during development into many more neuronal and non-neuronal (e.g. glia) cell types[46, 47]. Presumably, for this reason, these enhancers show relatively broad expression in the adult brain [45]. Subpallial enhancers as expected tended to drive expression in GABAergic cells, but do not distinguish between the various known subtypes of GABAergic interneurons[41, 48]. Therefore, although these tools are valuable to the elucidation of cell lineages, they are not necessarily more specific than promoter-based transgenic lines[15], which as noted earlier are not always specific enough for the analysis of native neural circuits.

Thus, a seemingly trivial difference in technique results in a large increase in utility for systems neuroscience. Applying the same methods discussed above to microdissected adult cortical subregions allows one to make molecular genetic tools apparently specific to particular cell types of the targeted brain regions. The microdissection is not in fact a trivial feature: by examining four subregions of the cortex separately, we found around four times as many reproducible peak calls as was obtained from the entire cortex[27], even though these four subregions together comprise a small minority of the cortex. This implies that individual cortical subregions contain their own epigenetically distinct cell types, which are washed out when pooled. Similarly, there is relatively little overlap between the enhancers active in the embryonic brain and those we have obtained from adult brain (Figure 2). Hence, it would be interesting to work backwards and study the developmental expression of EDGE lines made from subdivisions of the adult brain to investigate the genetic signatures of the pre-and postnatal processes that specify the enormous variety of neuronal cell types present in the fully differentiated adult brain. In sum, we do not claim to have discovered anything novel about transcription in the brain, although the sheer number of novel putative enhancers unique to particular cortical subregions was surprising, nor the methods described herein. What we claim is both novel and significant is the application of these methods to the generation of anatomically-specific tools enabling the study of the circuit dynamics of the adult brain[49].

It is worth mentioning that EDGE is not the same as enhancer traps. In enhancer traps[12, 30], one randomly inserts a minimal promoter construct into the genome in the hopes of integrating near a specific enhancer while EDGE involves the identification and use of enhancers specific to particular brain regions. The key advantage of EDGE over enhancer traps is anatomical targeting. To illustrate, we can compare our results to those of a recently published enhancer trap study[12] using a lentiviral vector containing the exact same minimal promoter we used. Since we are interested in the EC, we consider the creation of EC specific lines the goal, as in the current study. The total number of genotypically positive founders that express in the brain are similar (45/105: 43% herein vs. 42/151: 28%), and both techniques can yield very specific expression patterns. The key difference is the numbers of lines expressing in the EC at all (41/45: 91% herein vs. 6/42: 14%) and especially those more or less specifically expressing in the EC (16/45: 36% vs. 0/42: 0%). This illustrates the difference between the two approaches: enhancer traps result in expression in random cell types throughout the brain (and indeed the entire body), while EDGE targets those cell types found in particular brain regions of interest.

Of course, not everyone is interested in the EC. While we subtracted out any enhancers which expressed anywhere but the MEC (or LEC), other investigators interested in other brain regions can use the same strategy to develop tools specifically targeting their brain regions of interest. This process can occur for any brain regions, potentially providing cell type-specific tools to interrogate any neural circuit. Moreover, the more subdivisions of the brain one collects, the more one can subtract, so therefore the more specific the resulting putative enhancers will be. With this in mind we have initiated a second round of enhancer ChIP-seq with over 20 brain subregions which will provide a much more generally useful resource. Finally, the relatively small size of the enhancers means they can fit easily in viral vectors. If EDGE viruses recapitulate the anatomical specificity seen in transgenic mice, this will potentially bring EDGE to bear on any species[41]. This could revolutionize not only systems neuroscience, but ultimately provide a novel therapeutic avenue to rectify the circuit imbalances that underlie disorders of the central nervous system.

### Do enhancers specify neuronal cell types in the brain?

One of the most interesting questions in neuroscience is how we should think about the 100 or so billion neurons in our brains-as unique actors, or as repeated elements in a printed circuit? The answer is likely in between. Several investigators have proposed a canonical circuit for the neocortex[50, 51], with regional variations, and there are clearly commonalities in neocortical circuits, particularly with regards to layer-specific connectivity. Yet, within this general canonical theme there are uniquely specialized cell types in individual cortical subregions. Our results demonstrate that there are thousands of putative enhancers unique to cortical subregions, a number far larger than the number of genes that are specific to these subregions (indeed to our knowledge there are no EC specific genes). Why do the same genes use different enhancers to express in different cortical subregions? There is not yet enough data for a satisfactory answer, but the developmental literature discussed above would suggest a combinatorial code of transcription factors and active enhancers for each unique cell fate. If so, enhancer usage could provide a finer grained differentiation of cell type than gene expression alone. The fact that there are hundreds to thousands of unique enhancers in individual cortical subregions means that the genetic machinery exists to have a similar number of differentiable cell types. In support of this, a recent study of the transcriptome of thousands of individually sequenced neurons from two different cortical regions finds a large number of distinct transcriptional profiles between excitatory, but not inhibitory neurons[52]. This (as well as the fact that inhibitory neurons are a small minority of cortical neurons) may explain why we only obtained expression in excitatory neurons when we selected region-specific enhancers.

EDGE allows the generation of tools that provide a means to investigate the nature of neuronal cell types. For example, three of the enhancer constructs presented here drive expression in layer II of MEC, two of which (MEC-13-53 and MEC-13-81) exclusively in reelin-positive neurons (Figure 5 and data not shown for MEC-13-81). MEC LII reelin-positive neurons are stellate cells, which is arguably a cell type, but neither line expresses in 100% of reelin-positive neurons. There are two possible explanations for this is the first one being that these distinct enhancers drive expression in functionally distinct subsets of stellate cells[52, 53]. The other, possibility is that each enhancer drives expression in stellate cells as part of a co-regulated network of enhancers specifying this cell type[38]. If so, the difference in percentage of expression in stellate cells is largely artefactual, resulting from differential penetrance of transgene expression of otherwise identical cells due to mosaicism arising from insertional effects. The exhaustive biochemical, anatomical and electrophysiological characterization of each line necessary to provide a definitive answer to the relationship between these enhancers and cell types is beyond the scope of this paper. However, the fact that there are so many enhancers unique to specific cortical subregions implies the potential for a more direct connection between the molecular identity of a cell and cell type in other terms. Moreover, it is entirely possible that further subdivisions of the cells specified by these transgenic lines could provide even more specific expression. This could be achieved in a variety of ways, for example by finer manual microdissection, laser capture microscopy or even nested ChIP-seq of transgenically-labeled cells isolated by a cell sorter from microdissected tissue.

Regardless, our results certainly do not suggest that every enhancer defines a distinct cell type, in fact several of our lines express in more than one layer. There is not necessarily a one-to-one correspondence between cell types and enhancers: a single cell type could be specified by multiple unique enhancers, i.e. a co-regulated enhancer network[38]. Conversely, different cell types may arise from distinct combinatorial codes of active enhancers, meaning the number of different cell types may conceivably be even larger than the number of unique enhancers. Finally, there are other reasons for differential sets of active enhancers beyond definition of cell type: neural activity changes the chromatin landscape[54]. This means that activity of differential enhancers does not automatically imply different cell types but changes in function of a given cell. Nevertheless, differential enhancer utilization does signify distinct epigenetic signatures, even if their functional significance is currently unclear. We therefore maintain that a powerful way to investigate the relationship of diverse cis-acting elements of the genome to the functional circuitry of the brain is to create and study enhancer-specific tools like those presented here.

## Acknowledgements

This work was supported by the FRIPRO ToppForsk grant Enhanced Transgenics (90096000) of the Research Council of Norway, the Kavli Foundation, the Centre of Excellence scheme of the Research Council of Norway – Centre for Biology of Memory and Centre for Neural Computation, The Egil and Pauline Braathen and Fred Kavli Centre for Cortical Microcircuits and the National Infrastructure scheme of the Research Council of Norway – NORBRAIN. We would like to thank Haiyan Wu and Qiangwei Zhang for their help with *In Situ hybridization*, Ute Hostick of the transgenic mouse facility in Eugene, Oregon for her help with pronuclear injection and Hanne Mali MøllergÅrd for her help with genotyping.

## Author contribution

Conceptualization: J.C. and C.K.; Software: J.C.; Validation: S.B.; Methodology, Formal analysis, Data curation, and Visualization: J.C. and S.B.; Investigation and Resources: S.B., M.P.W. and J.C.; Writing – original draft: S.B., J.C. and C.K.; Writing – review and editing: S.B., M.P.W., J.C. and C.K.; Supervision: J.C., J.N. and C.K.; Project administration: J.N. and C.K.; Funding acquisition: C.K.

## METHODS

### Contact for reagent and resource sharing

Further information and requests for resources and reagents should be directed to and will be fulfilled by the Lead Contact, Cliff Kentros.

### Experimental model and subject details

The experimental model used in this study is the rodent *M. musculus*. For the microdissection two C57BL/6J mice (Jackson Laboratory, stock no. 000664, P56, one male, one female) were used. For pro-nuclear injection B6D2F1 mice were used. The resulting offspring were backcrossed with C57BL/6J mice and various reporter lines. The mice used for histology were a mix of male and female, taken from several litters when multiple individual mice were investigated, and were at least 6 weeks old.

All mice were kept on a 12-h light/12-h dark schedule in a humidity-and temperature-controlled environment. All experiments in Norway were performed in accordance with the Norwegian Animal Welfare Act and the European Convention for the Protection of Vertebrate Animals used for Experimental and Other Scientific Purposes. All experiments involving animals in Oregon (pronuclear injection and husbandry of the resulting animals) were performed in accordance with guidelines approved by University of Oregon’s Animal Care and Use Committee and the National Institutes of Health Guide for the Care and Use of Laboratory Animals (National Institutes of Health Publications No. 80-23).

### Method details

#### Microdissection

Two C57BL/6J mice (P56) were deeply anesthetized by injection with pentobarbital (100mg/ml in 96% ethanol, Ås produksjonslab AS). The brains were removed and horizontal or coronal 500 µm sections were cut on a Leica VT 1000 S microtome and kept at 4 °C until dissection. Bilateral dissection was performed, while watching the tissue through a dissection microscope with transmitted and reflected white light (Zeiss Discovery V8 stereomicroscope) applying architectonic criteria [55-59] to unstained tissue. The tissue samples were snap-frozen in liquid nitrogen, kept at -80^0^C and shipped on dry ice.

All dissections avoided border regions (i.e., centered in the identified cortical area). In horizontal sections, MEC is easily recognized by the marked shape of the cortex, the prominent white, opaque lamina dissecans and the radial organization of the layers deep to the lamina dissecans. Layer II neurons are large spherical neurons, which differ markedly in level of opacity from those in layer III. The medial border between MEC and parasubiculum is characterized by the loss of the differentiation between layers II and III, and the border with the laterally adjacent postrhinal cortex is characterized by the loss of the large spherical neurons in layer II [55-59]. We only sampled the more dorsal and central portions of MEC. LEC shares the large layer II neurons with MEC, but the radial organization in layer V is absent. The anterior and dorsal border of LEC with the perirhinal cortex is characterized by the abrupt disappearance of the large layer II neurons. We only sampled the most lateral portions of LEC, as to avoid contamination with ventromedially adjacent components of the amygdaloid complex. ACC and RSC were sampled from the medial wall of the lateral hemisphere above the corpus callosum, avoiding the most anterior part of ACC and the posteroventral part of RCS. Since the border between the two areas coincides with the dorsal-anterior tip of the hippocampal formation, all samples avoided that border region.

In coronal sections, ACC and RSC samples were taken dorsal to the corpus callosum, just below the shoulder of the medial wall of the hemisphere down to, but not touching the corpus callosum, as to avoid inclusion of the indusium griseum. Samples were taken from sections anterior to the most anterodorsal tip of the hippocampal formation in case of ACC and posterior to the tip in case of RSC. Samples of LEC were collected one section after the disappearance of the piriform cortex characterized by a densely packed thick layer II, a polymorph lightly packed deeper cell layer and the presence of the endopiriform nucleus. LEC shows cytoarchitectonic features similar to those described above. We sampled only from the vertical part of LEC, directly below the rhinal fissure. For MEC, samples were collected from more posterior coronal sections, using shape of the section, the presence of the ventral hippocampus and cytoarchitectonic features as described above as our selection criteria.

#### ChIP-seq

All dissected brain tissues were briefly homogenized and cross-linked with 1% formaldehyde at room temperature with rotation for 15 min. Cross-linking was quenched with glycine (150mM in PBS), then tissue was washed and flash frozen. Chromatin was extracted as previously described [25, 60]. Briefly, nuclei were extracted, lysed and sonicated (30 min, 10-sec pulses) to produce sheared chromatin with an average length of ∼250 bp. 1-10µg of final soluble chromatin was used for each ChIP and combined with Protein G Dynabeads (Invitrogen, cat# 10004D) prebound with 5µg of antibodies to H3K4me2 (Abcam ab7766) or H3K27ac (Abcam ab4729). Immunoprecipitated chromatin was washed five times with 1mL of wash buffer and once with TE. Immunoprecipiated chromatin was eluted, cross-links were reversed, and DNA was purified. Libraries were prepared for sequencing using NEBNext ChIP-Seq Library Prep reagents and sequenced on the Illumina HiSeq 2000 platform at the Yale Center for Genome Analysis.

#### Cloning of transgenic constructs

The putative enhancers sequences were cloned from BACs (chori.org) and transferred to pENTR^tm^/D-TOPO^®^ vectors by TOPO^®^ cloning (Invitrogen, K2400-20). The putative enhancers were transferred to injection plasmids by gateway cloning^®^ (Invitrogen, 11791-019). The resulting plasmids consist of a putative enhancer followed by a mutated heatshock promoter 68 (HSP68), a tTA gene, a synthetic intron and a WPRE element (Figure S3).

#### Pronuclear injection

The 10 injection plasmids were linearized by enzyme digestion to keep the relevant elements but remove the bacterial elements of the plasmids. Linearized vectors were run on a 1% agarose gel and isolated using a Zymoclean Gel DNA Recovery Kit (Zymo research, D4001). Fertilized eggcells were injected with 1 µl of DNA at concentrations of 0.5 to 1 ng/µl, leading to surviving pups of which 96 were genotypically positive for MEC and 9 were genotypically positive for LEC (Table S1). Pronuclear injections were done at the transgenic mouse facility of the University of Oregon.

#### Mouse husbandry

Mouse lines were named after the ranked enhancers identified in this study, as specified in Data S1. The nomenclature consists of firstly the targeted region, secondly the year of microdissection, thirdly the rank of the enhancer that corresponds with the row in Data S1 and finally a letter for the founder. To illustrate, line MEC-13-53A is based on MEC tissue isolated in 2013, where the particular enhancer was ranked 53 and the founder is specified by the “A”.

All genotypically positive founders based on MEC enhancers were initially mated with histone GFP mice (Jackson laboratory, Tg(tetO-HIST1H2BJ/GFP)47Efu, stocknr. 005104), while those based on LEC enhancers were mated with GCaMP6 mice (in house made). Double positive pups were used for further analysis. Subsequent crosses were done with GCaMP6 mice (in house made), TVAG mice (Line TVAG5 from [61]), ArChT mice [31], tetO-eGFP (Jackson laboratory, C57BL/6J-Tg(tetO-EGFP/Rpl10a)5aReij/J_JAX) and hM3 mice [8].

#### Genotyping

Genotyping was done on ear tissue using a Kapa mouse genotyping kit (Kapa Biosystems, Cat# KK7302). Primer pairs for the appropriate gene and internal controls (Table S2) are added to the PCR mixture at a final concentration of 10µM. The PCR reaction was done by an initial step of 4 minutes at 95°C, then 20 cycles of 1 minute at 95°C, 30 seconds at 70°C reduced by 0.5°C each cycle, and 30 seconds at 72°C. This is followed by 20 cycles of 30 seconds at 95°C, 30 seconds at 60°C, and 30 seconds at 72°C and a final 7 minute step at 72°C. The products are run on a 1% agarose gel along with positive and negative controls.

#### In situ hybridization

Double positive mice (tTA+/-, reporter gene+/-) were deeply anesthetized with pentobarbital and transcardially perfused with 0.9% saline first and freshly made 4% formaldehyde (in 1x DPBS, thermofisher, Cat# 14200075) second. At least 2 mice from different litters were investigated on consistent expression patterns. Brains were removed and postfixated overnight in 4% paraformaldehyde. Subsequently the brains were dehydrated for at least 24h with 30% sucrose in 1x PBS. The brains were sectioned sagittally at 30µm on a Cryostat, mounted directly (on Fisherbrand Superfrost Plus microscope slides (Fisher Scientific Cat #12-550-15)) and dried overnight at room temperature. Slides were stored at -80°C.

Slides were thawed in closed containers. Sections were outlined with a PAP pen (Sigma, cat# Z377821-1EA). The probe was diluted (usually 0.1-1µgm/ml) in hybridisation buffer (1:10 10x salt solution, 50% deionized formamide (sigma,cat# D-4551), 10% dextran sulfate (sigma, cat# D-8906), 1mg/ml rRNA (sigma, Cat#R5636), 1x Denhardt’s (Sigma cat# D-2532). Salt solution (10x) was made with 114g NaCl, 14.04g TrisHCl, 1.3g TrisBase, 7.8g NaH2PO4.2H2O, 7.1g Na2HPO4 in H_2_O to 1000ml with a final concentration of 0.5M EDTA). The probe was denatured for 10 min at 62°C, added to the section and coverslipped (Fisher, cat# 12-548-5P). The slides were incubated overnight at 62°C in a closed box with filter paper wetted in 1x SSC with 50% formamide.

The slides were transferred to polypropylene Coplin jars containing 1x SSC with 50% formamide and 0.1% Tween-20 warmed to 62°C for 10 minutes to allow the coverslips to fall off. The slides were washed 3×30 minutes at 62°C. Then the slides were washed 3×30 minutes in MABT (11.6g Maleic acid (sigma, cat#M0375-1kg), 8.76g NaCl, 5ml 20% tween, pH 7.5, ddH_2_O to 1000ml) at room temperature.

The slides were drained (not dried) and re-circled with a PAP pen. Then blocking solution was added (600µl MABT, 200µl sheep serum, 200µl 10% blocking reagent (Roche cat#11 096 176 001)) and slides were incubated in a Perspex box with wetted filter paper at room temperature for 2-3 hours. The slides were drained and 1:5,000 sheep anti-dig AP in blocking solution was added followed by overnight incubation.

4g of polyvinyl alcohol was dissolved into 40ml AP staining buffer (100mM NaCl, 50mM MgCl_2_, 100mM Tris pH9.5, 0.1% Tween-20) by heat and cooled to 37°C. The slides were washed in MABT 5 times for 4 minutes. And subsequently washed 2×10 minutes in AP staining buffer. Nitroblue tetrazolium chloride (Roche, cat# 11 383 213 001. At 3.5 µl/ml), 5-Bromo-4-chloro-3-indolyl-phosphate,4-toluidene salt (Roche, cat# 11 383 221 001. At 2.6 µl/ml) and Levamisole (Vector, cat# SP-5000. At 80µl/ml) was added to the cool polyvinyl alcohol solution. This was shaken well and transferred to a Coplin jar. The slides were added to the jar and incubated at 37°C for 3 to 5 hours. The reaction was stopped by washing in 2xPBS with 0.1% Tween-20. The slides were subsequently wash 2X in ddH2O, and dehydrated quickly through graded ethanols from 50%, 70%, 95% to 100% ethanol. Finally the slides were cleared in xylene and coverslipped.

#### Immunohistochemistry

Double positive mice (tTA+/-, TVAG+/-or eGFP +/-) were deeply anesthetized with pentobarbital and transcardially perfused with approximately 30ml 0.9% saline first and approximately 30ml freshly made 4% paraformaldehyde (in 1x DPBS, thermofisher, Cat# 14200075) second. Brains were removed and postfixated for 24 hours in 4% paraformaldehyde. Subsequently the brains were dehydrated with 30% sucrose in 1x PBS. The brains were sectioned horizontally at 50µm and kept in TCS (tissue collection solution, 25% glycerol, 35% ethyl glycol, 50% 1xDPBS) at -20°C.

Immunohistochemistry was done by two initial 10 minute washes in 1xDPBS and subsequent permeabilized by a 60 minute wash in 1% Triton X-100 (Sigma, Cat#T9284) in 1xDPBS. Then the tissue is incubated in primary antibody in 1xDPBS with 1% trition X-100 and 5% donkey serum (Sigma, Cat# D9663) for 48 hours at 4°C. Primary antibodies and dilutions were: Rabbit-anti-2A (1:2000, Millipore, cat#ABS31), Mouse-anti-reelin (1:1000, Millipore, cat# Mab5364), Mouse-anti-GAD67 (1:1000, Millipore, cat# Mab5406), Mouse-anti-calbindin (1:10000, Swant, cat# CB300).

After incubation with primary antibodies, sections were washed 4x in 1xDPBS (10 minutes per wash) and 2x in 1xDPBS with 1% Triton X-100. Then sections were incubated for 6h at room temperature in secondary antibody (all secondary antibodies were raised in Donkey and diluted 1:250). The secondary antibodies were: anti-Rabbit-AF488 (Jackson ImmunoResearch, Cat# 711-545-152) and anti-Mouse-Cy^tm^3 (Jackson ImmunoResearch, Cat# 715-165-151)

The sections were DAPI stained by a single 10 minute wash in 1xDPBS with 0.2µg/ml DAPI (thermofisher, D1306) and finally washed 5x (10 minutes per wash) in 1x DPBS. Sections were mounted on superfrost^®^ plus glass slides (VWR, Cat# 631-9483) and coverslipped with polyvinyl alcohol with 2.5% DABCO (Sigma, Cat# D27802).

#### Imaging

From mice in the lines MEC-13-53A x TVAG and MEC-13-104B x tetO-eGFP MEC was imaged in sections from three different dorsal-ventral levels with a Zeiss Meta 880 confocal microscope. Two channels were imaged: one for AF488 with maximum excitation wavelength at 488nm and maximum emission wavelength at 528nm and one for Cy3 with maximum excitation wavelength at 561nm and maximum emission wavelength at 595nm.

For display images, sections were imaged on Zeiss Axio.scan Z1 scanners in three preset channels: DAPI, dl488 and dl549.

#### Image processing

From the Zeiss proprietary file format .lsm, .tiff files were exported. These were processed in Adobe Photoshop and all alterations in levels were made on the entire images. In some cases, images were processed to remove visual artifacts and background.

### Quantification and statistical analysis

#### ChIP-seq data analysis

ChIP-Seq data was initially processed as previously described (Reilly et al 2015). Briefly, reads were aligned to the mm9 version of the mouse genome using bowtie (v1.1.1) [62]. Enriched regions were identified in individual replicates using a sliding window method as previously described [63]. Enriched regions were divided into functional categories based on overlaps with genomic features as annotated by Ensembl v67 using Bedtools (2.19.0) [64]. Reproducibly enriched regions were determined as the union of overlapping regions identified in both biological replicates. Putative enhancer regions from intergenic and intronic portions of the genome were then assigned target genes using GREAT. H3K27ac ChIP-Seq reads were retrieved from Encodeproject.org for 17 mouse tissues [27] and uniformly processed as above. Enhancers for all cell types were combined and merged to generate a uniform annotation of all possible enhancers. H3K27ac counts at each enhancer from each tissue were calculated using mrfQuantifier [65]. Pearson correlations for all enhancer signals were calculated and plotted using R (https://www.r-project.org/)K-means clustering of H3K27ac count matrix was performed using Cluster (v3.0) [66]. Rows were centered on the mean value of the row and normalized, the k parameter was the total number of tissues, and 100 runs were performed. The clustering result was then visualized using Java TreeView [67]. Subregion specific clusters of enhancers were intersected with peak calls from all other tissues to identify enhancers with likely tissue specific function. Subregion specific enhancers were assigned two target genes using GREAT, ranked by H3K27ac signal, and overlapped with vertebrate conserved sequences [68].

#### Counting

Mice in the lines MEC-13-53A x TVAG and MEC-13-104B x tetO-eGFP MEC was imaged on sections from three different dorsal-ventral levels. For each strain 4 mice from at least 2 litters each were used. For each section, three to seven slices in the Z direction with 1.5µm spacing were taken, with a 20x objective and tiling to cover the entire MEC. Counts were made on the confocal images for single positive cells expressing transgenes, cells expressing native genes (GAD67, Reelin, Calbindin) and cells expressing both. Graphs were made in Microsoft Excel and statistical analysis was done in SPSS (v22, IBM).

### Data and software availability

No new software was generated in this study.

Raw data can be accessed on GEO under accession number GSE112897.

Processed data can be viewed here.

**Supplemental Information:** Figures S1-S7 and Tables S1 and S2

**Data S1:** Ranked regionally specific enhancers for brain regions MEC, LEC, ACC and RSC, related to Figure 2.

## KEY RESOURCES TABLE

| REAGENT or RESOURCE | SOURCE | IDENTIFIER |
| --- | --- | --- |
| **Antibodies** |  |  |
| H3K4me2 | Abcam | Cat# ab7766, RRID:AB_2560996 |
| H3K27ac | Abcam | Cat# ab4729, RRID:AB_2118291 |
| Rabbit-anti-2A | Millipore | Cat# ABS31, RRID:AB_11214282 |
| Mouse-anti-reelin | Millipore | Cat# Mab5364, RRID:AB_2179313 |
| Mouse-anti-GAD67 | Millipore | Cat# Mab5406, RRID:AB_2278725 |
| Mouse-anti-calbindin | Swant | Cat# CB300, RRID:AB_10000347 |
| anti-Rabbit-AF488 | Jackson ImmunoResearch | Cat# 711-545-152, RRID:AB_2313584 |
| anti-Mouse-Cy<sup>tm</sup>3 | Jackson ImmunoResearch | Cat# 715-165-151, RRID:AB_2315777 |
| **Biological Samples** |  |  |
| Healthy, adult microdissected brain tissue (mouse) | This paper | N/A |
| **Critical Commercial Assays** |  |  |
| NEBNext ChIP-Seq Library Prep | NEB | E6240L |
| **Deposited Data** |  |  |
| Raw data | GEO | GSE112897 |
| Processed data | NCBI session | http://genome.ucsc.edu/cgi-bin/hgTracks?hgS_doOtherUser=submitt&hgS_otherUserName=Jcotney&hgS_otherUserSessionName=Kavli |
| **Experimental Models: Organisms/Strains** |  |  |
| C57BL/6J mice | Jackson Laboratory | Stock no. 000664, RRID: IMSR_JAX:000664 |
| B6D2F1 mice | Jackson Laboratory | Stock no. 100006, RRID:IMSR_JAX:100006 |
| TetO-GCaMP6 mice | This paper | P8966 |
| TetO-histoneGFP mice Tg(tetO-HIST1H2BJ/GFP)47Efu | Jackson laboratory | Stock no. 005104, RRID:IMSR_JAX:005104 |
| TetO-TVAG mice | From [61] | Line TVAG5 |
| TetO-ArchT mice | From [31] | Line tetO-ArchT2 |
| tetO-eGFP mice C57BL/6J-Tg(tetO-EGFP/Rpl10a)5aReij/J_JAX | Jackson laboratory | Stock no. 024898, RRID:IMSR_JAX:024898 |
| TetO-HM3 mice | From [8] | Line TRE-hM3Dq, RRID:IMSR_JAX:014093 |
| MEC-13-32B | This paper | N/A |
| MEC-13-53A | This paper | N/A |
| MEC-13-53B | This paper | N/A |
| MEC-13-53D | This paper | N/A |
| MEC-13-53G | This paper | N/A |
| MEC-13-81A | This paper | N/A |
| MEC-13-104B | This paper | N/A |
| MEC-13-123A | This paper | N/A |
| MEC-13-123B | This paper | N/A |
| MEC-13-123C | This paper | N/A |
| LEC-13-8B | This paper | N/A |
| LEC-13-108A | This paper | N/A |
| Oligonucleotides |  |  |
| Genotyping primers, Table S2 | This paper | N/A |
| Recombinant DNA |  |  |
| Injection construct (Gateway destination vector) | This paper | pDest-HSP68-tTA-WPRE |
| Software and Algorithms |  |  |
| Bowtie (v1.1.1) | [62] | http://bowtie-bio.sourceforge.net/index.shtml |
| Bedtools (2.19.0) | [64] | http://bedtools.readthedocs.io/en/latest/ |
| mrfQuantifier | [65] | http://archive.gersteinlab.org/proj/rna-seq/rseqtools/ |
| GREAT | [28] | http://great.stanford.edu/public/html/ |
| R | R-project | https://www.r-project.org/ |
| Cluster (v3.0) | [66] | http://bonsai.hgc.jp/~mdehoon/software/cluster/software.htm |
| Java TreeView | [67] | http://jtreeview.sourceforge.net/ |
| SPSS statistics v. 22 | IBM | https://www.ibm.com/products/spss-statistics |
| Creative Cloud (photoshop, illustrator) 2017 edition | Adobe | https://www.adobe.com/no/creativecloud.html |
| Excel 2016 | Microsoft | www.microsoft.com |
| Other |  |  |
| Illumina HiSeq 2000 | Illumina | N/A |

**Figure S1.**
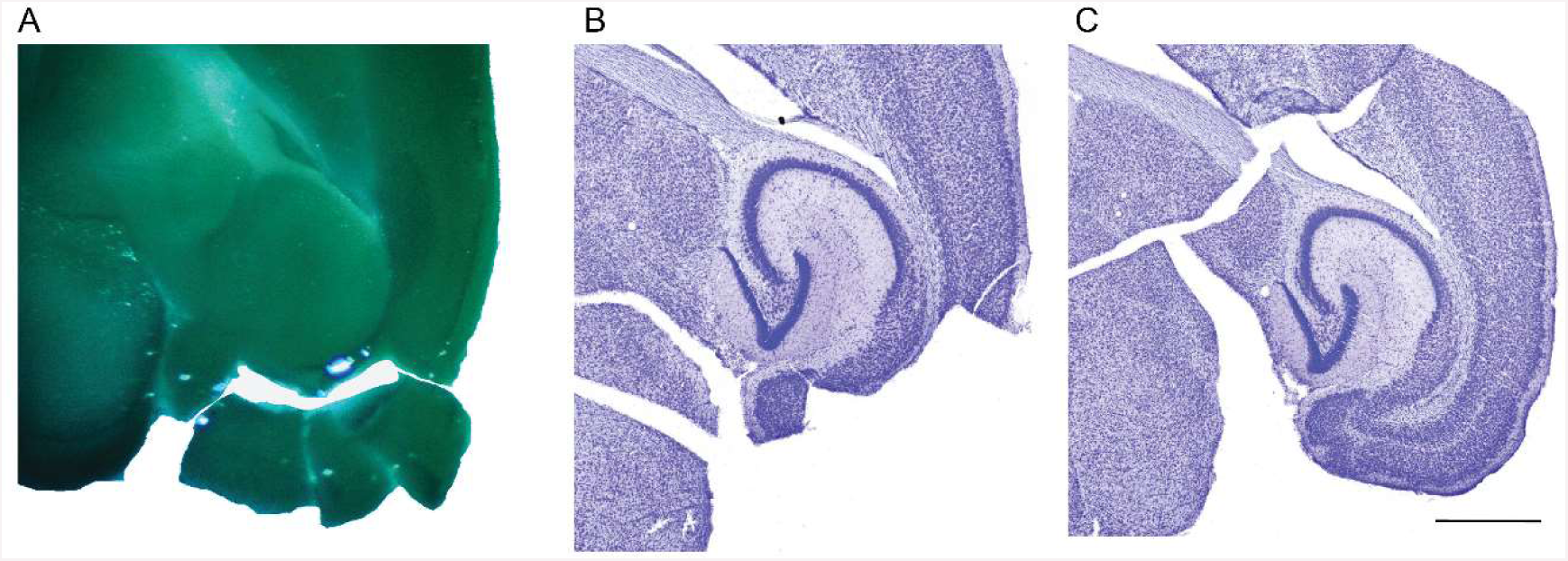
Example of microdissection, related to STAR methods and Figure 1. (A) 500µm thick section during microdissection. (B) Contralateral side of the same section as in (A), re-sectioned to 50µm and Nissl stained. (C) Re-sectioned (50µm), Nissl stained tissue from (A). Scalebar is 1000µm.

**Figure S2.**
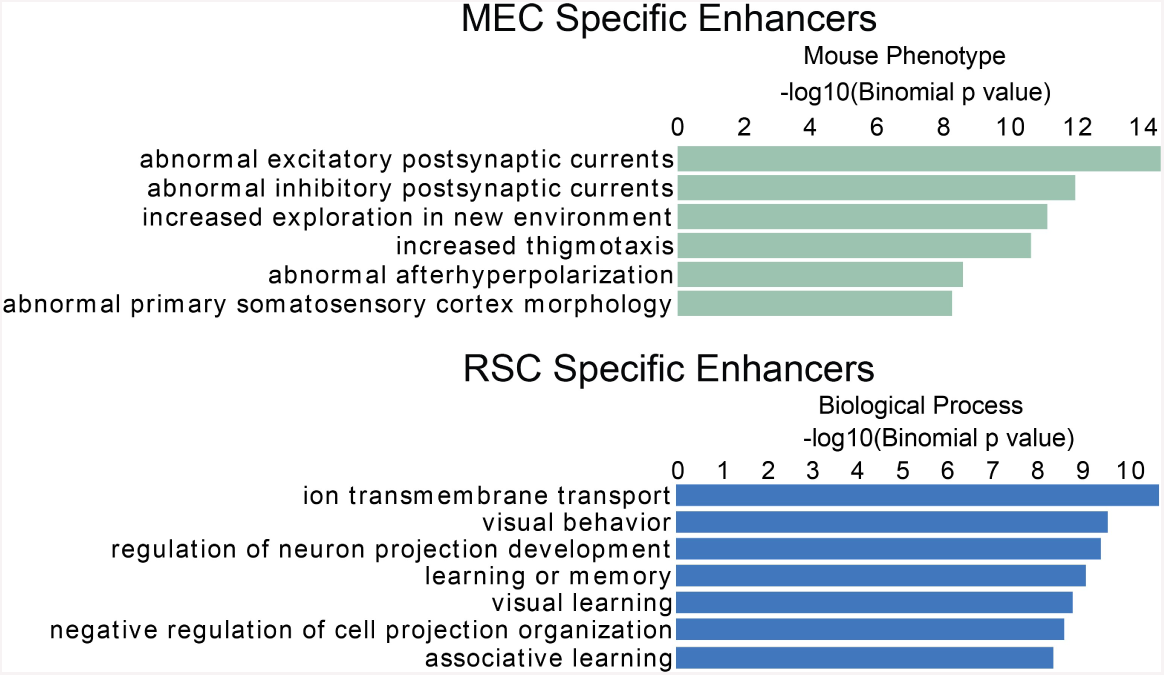
Gene Ontology of selected cortical subregions, related to Figure 2. Peak calls for each tissue type were used in the GREAT algorithm to assign genes. Ontology terms for these genes were ranked on Binominal P-value.

**Figure S3.**
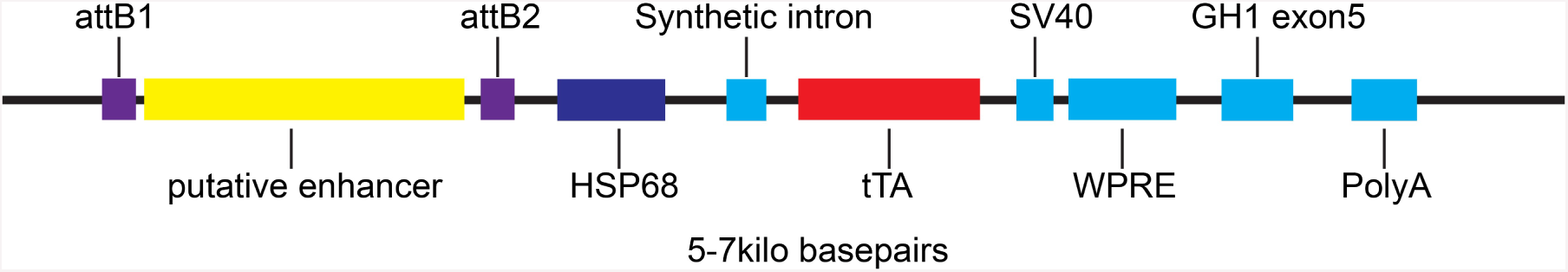
Injection construct, related to Figure 3. The putative enhancer of 0.7 to 3kbp was cloned to the injection construct by gateway^®^ cloning. The synthetic intron, SV40, WPRE and growth hormone 1 exon 5 are present for optimal mRNA stability and expression of the tetracycline TransActivator (tTA). The construct is linearized with appropriate restriction enzymes depending on exact sequence of the putative enhancer.

**Figure S4.**
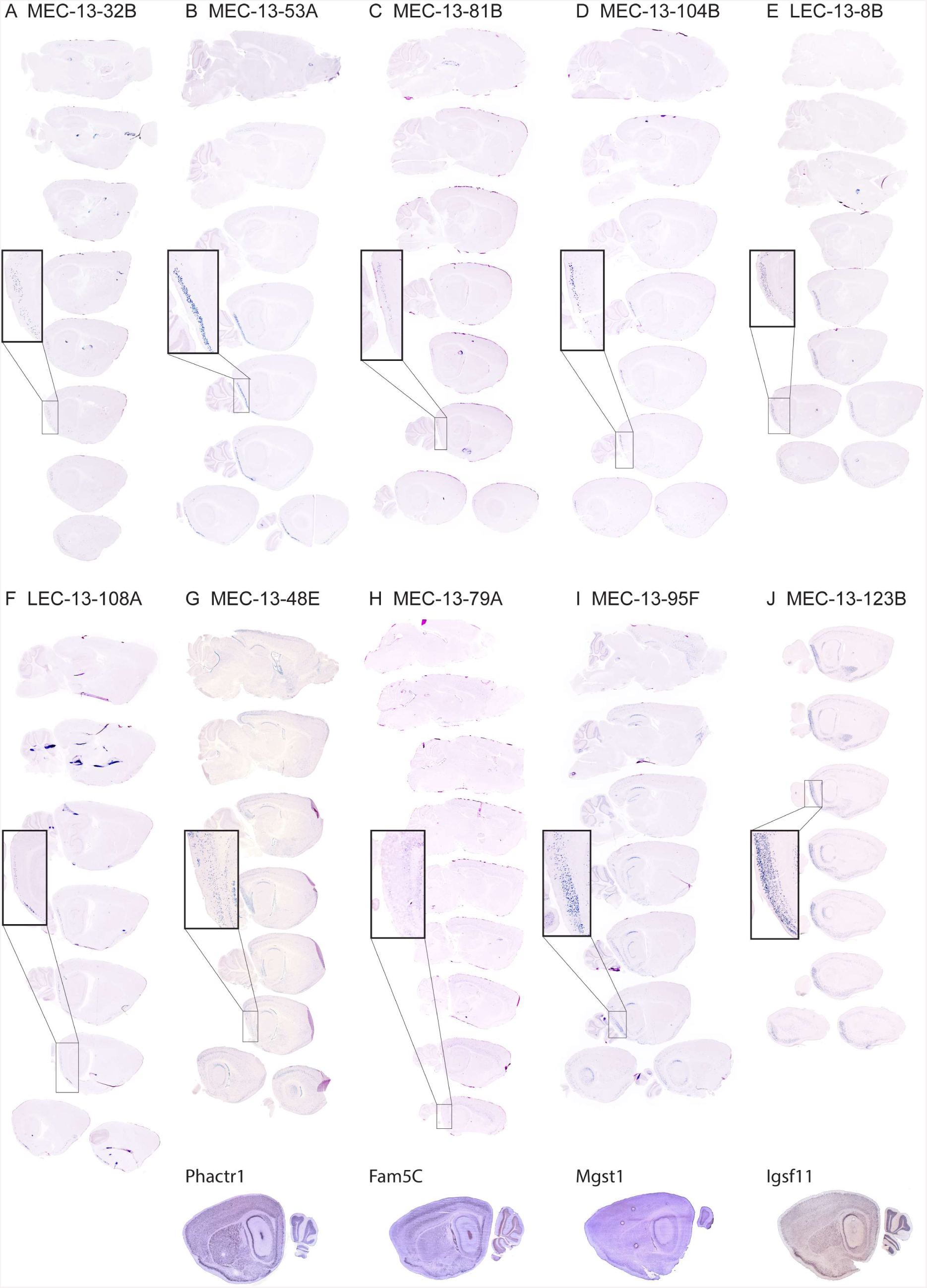
Extended medial lateral coverage of sagittal sections, related to Figure 3 and 4. (A)-(F) Enhancer lines based 6 different enhancers used (MEC-13-32B, MEC-13-53A, MEC-13-81B, MEC-13-104B, LEC-13-8B, LEC-13-108A) show specific transgene expression in the EC. (G)-(J) Enhancer lines based on 4 different enhancers (MEC-13-48E, MEC-13-79A, MEC-13-95F, MEC-13-123B) show enriched, but not specific transgene expression in the EC. The bottom row shows in *situ hybridization* (taken from brain-map.org) of associated genes.

**Figure S5.**
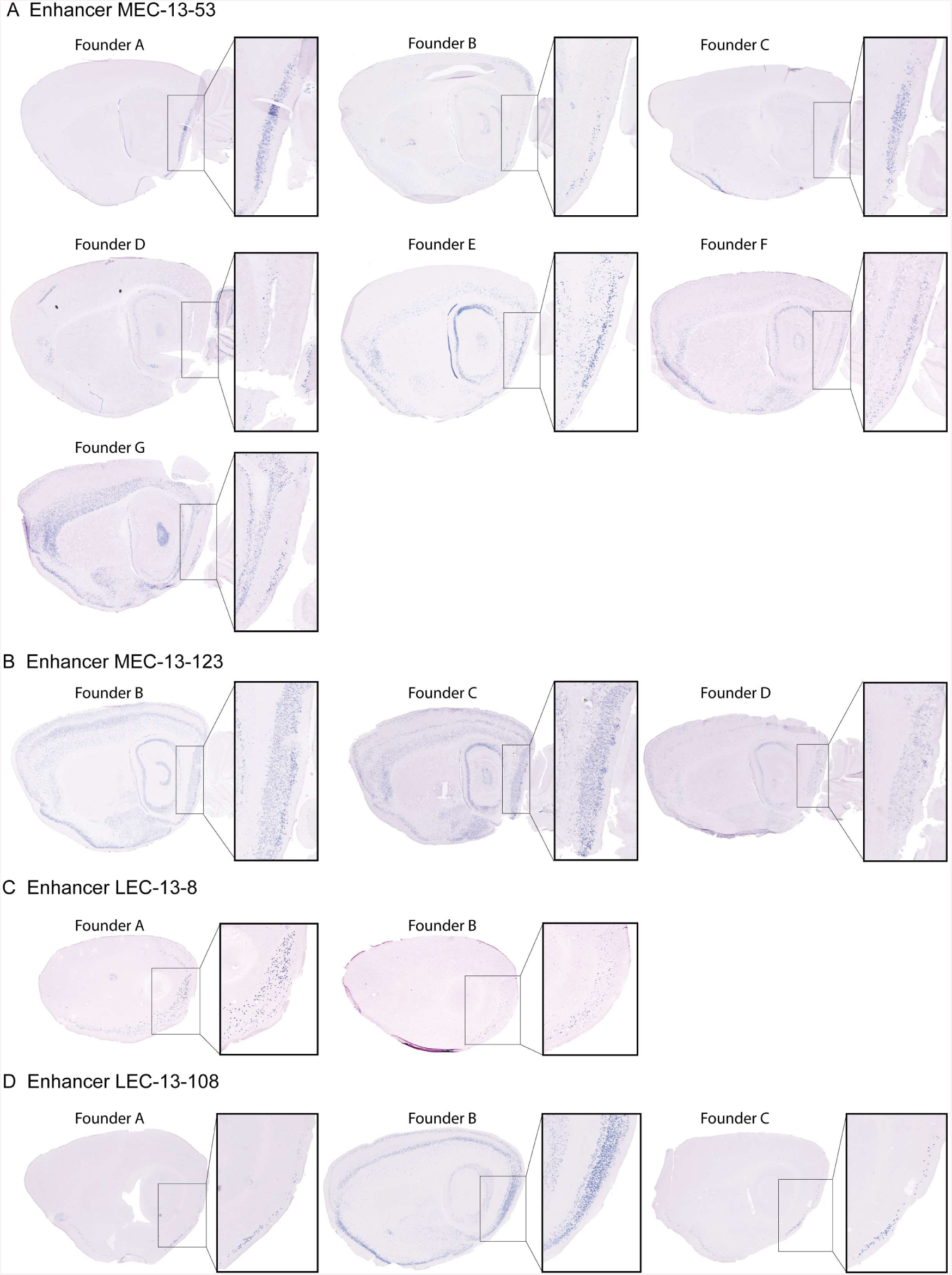
Enhancer driven transgene expression of various genomic insertions, related to Figure 4. Sagittal sections of approximately similar levels. Different founders based on the same enhancers show roughly similar expression patterns. All mice based on MEC specific enhancers were crosses with hGFP reporter mice, while all mice based on LEC specific enhancers were crossed with GC6 payload mice. (A) Enhancer MEC-13-53 reproducibly shows expression in LII of the EC in 6 of the 7 analyzed mouse lines. We do find expression in other regions, such as visual cortex in founder B, the CA fields of the hippocampus in founder E, and deep layers of cortex in founder G. But since all of these patterns of expression occur only once within the 7 analyzed lines, we consider them to be positional effects. (B) Enhancer MEC-13-123 reproducibly shows expression in LIII of the EC, CA3 and select cortical layers. (C) Enhancer LEC-13-8 reproducibly shows expression in LIII of the EC. (D) Enhancer LEC-13-108 reproducibly shows expression in LII of the EC. We consider the additional expression in the “founder B” line to be another positional effect.

**Figure S6.**
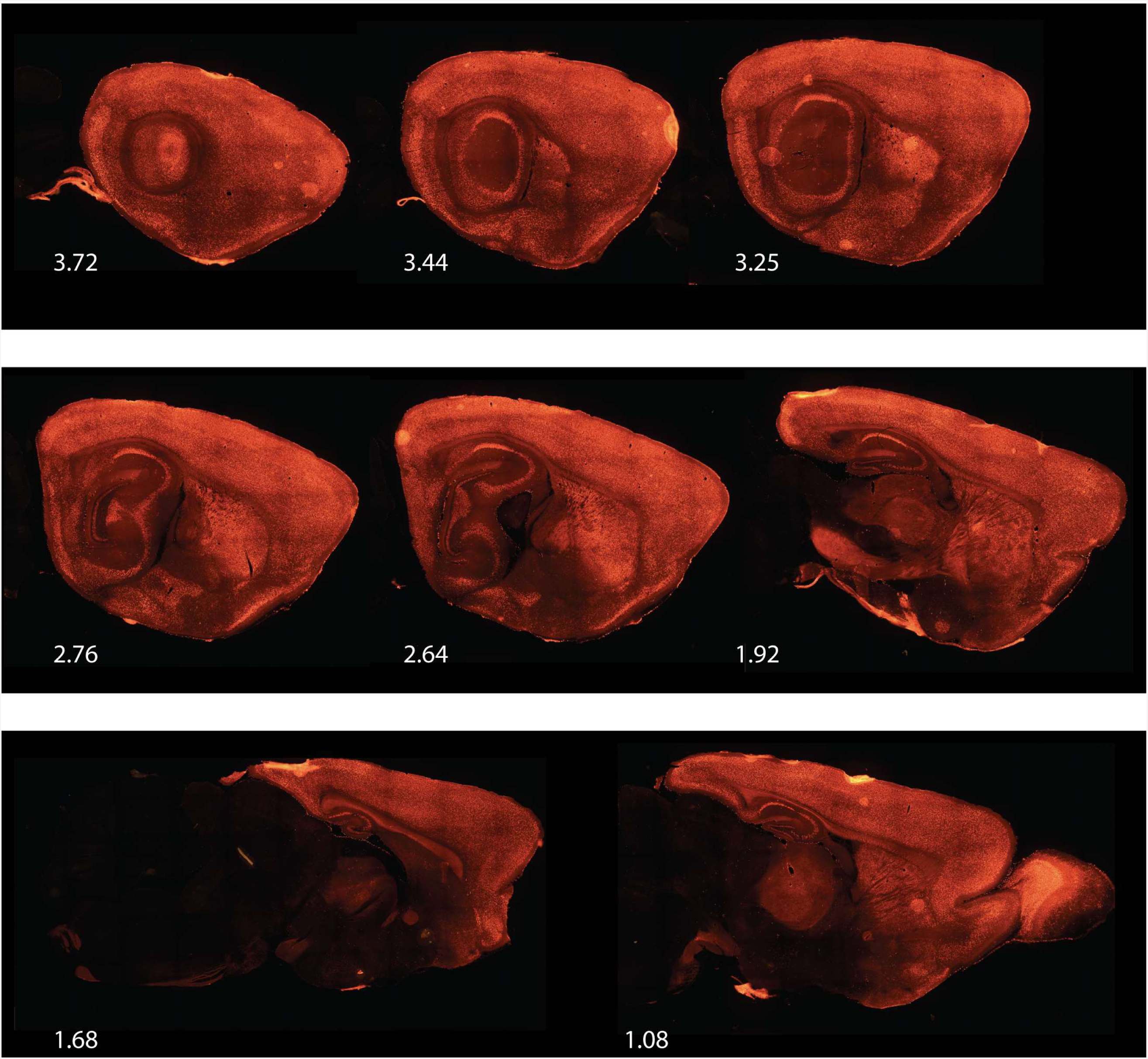
Expression of tTA dependent transgene gCaMP6 in a CaMKIIa driver line, related to main text describing Figures 3 and 4. Fluorescent signal of the mCherry conjugated to the tTA dependent gCaMP6 in sagittal sections (approximate level in mm lateral to midline indicated in white) show expression throughout the brain.

**Figure S7.**
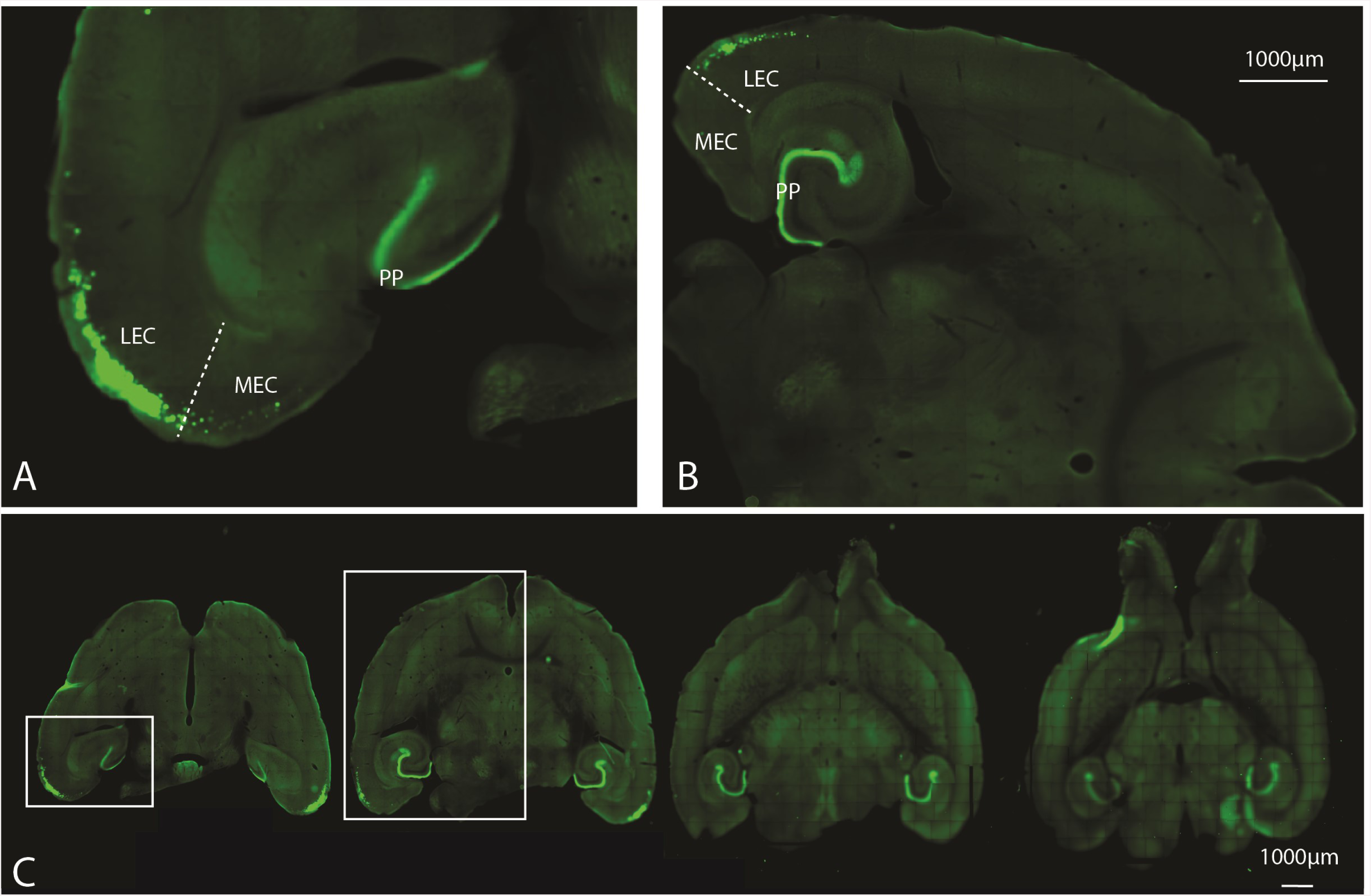
Expression of tTA dependent transgene gCaMP6 in enhancer line LEC-13-108A, related to main text describing Figures 5 and 6. The labeling of axons in the molecular layer in the dentate gyrus illustrates the transgenic labeling of layer II reelin+ cells.

**Table S1.** Overview of generated transgenic lines, related to Figures 3 and 4. Genomic coordinates are in mm9 and indicate the exact region used for the enhancer in the transgenic construct, rather than the putative enhancer as indicated by ChIP-seq. Enhancer rank indicates the rank as a result of the ChIP-seq analysis. Associated genes are a result of analysis of GREAT. All indications of expression are based on the initial round of assessment where mice based on MEC specific enhancers were crossed with histone GFP reporter mice and mice based on LEC specific enhancers were crossed with gCaMP6. The numbers in the last column are taken from Shima et al. 2016, table 1 and supplemental table 1 to identify the lines expressing in the EC (53L, 56L, TCAO, TCAR, TCIF, TCLC).

|  |  |  |  |  |  |  |  |  |  |  |  |  |  |
| --- | --- | --- | --- | --- | --- | --- | --- | --- | --- | --- | --- | --- | --- |
| Genomic coordinates enhancer | chr1:148,339,12<br>9-148,340,300 | chr16:39,750,78<br>9-39,753,454 | chr10:99,573,05<br>1-99,574,981 | chr15:50,913,89<br>6-50,916,356 | chr7:65,916,19<br>8-65,918,580 | chr6:138,334,72<br>8-138,335,952 | chr8:49,906,38<br>8-49,908,569 | chr13:42,782,50<br>3-42,784,035 | MEC | chr2:171,158,07<br>9-171,159,156 | chr5:118,194,6<br>53-118,195,333 | LEC | Shima et al. |
| Enhancer rank | 79 | 123 | 81 | 104 | 32 | 95 | 53 | 48 |  | 8 | 108 |  |  |
| Associated genes | Fam5c | Igsf11 | Kitl | Trps1 | Ube3a | Lmo3 | Odz3 | Phactr1 | total | Dok5 | Nos1 | total | variable |
|  |  |  | Gm4301 | Eif3h | Atp10a | Mgst1 |  | Tbc1d7 |  | Cbln4 | Ksr2 |  | integration |
| tTA positive founders | 6 | 6 | 10 | 12 | 8 | 16 | 18 | 20 | 96 | 5 | 4 | 9 | 151 |
| Lines analyzed | 5 | 4 | 7 | 8 | 4 | 13 | 16 | 16 | 73 | 4 | 3 | 7 | 151 |
| GFP signal in EC | 2 | 4 | 3 | 3 | 3 | 7 | 7 | 7 | 36 | 2 | 3 | 5 | 6 |
| No GFP signal in brain | 2 | 0 | 4 | 4 | 1 | 5 | 9 | 8 | 33 | 2 | 0 | 2 | 109 |
| GFP in brain but no GFP in EC | 1 | 0 | 0 | 1 | 0 | 1 | 0 | 1 | 4 | 0 | 0 | 0 | 36 |

**Table S2.**
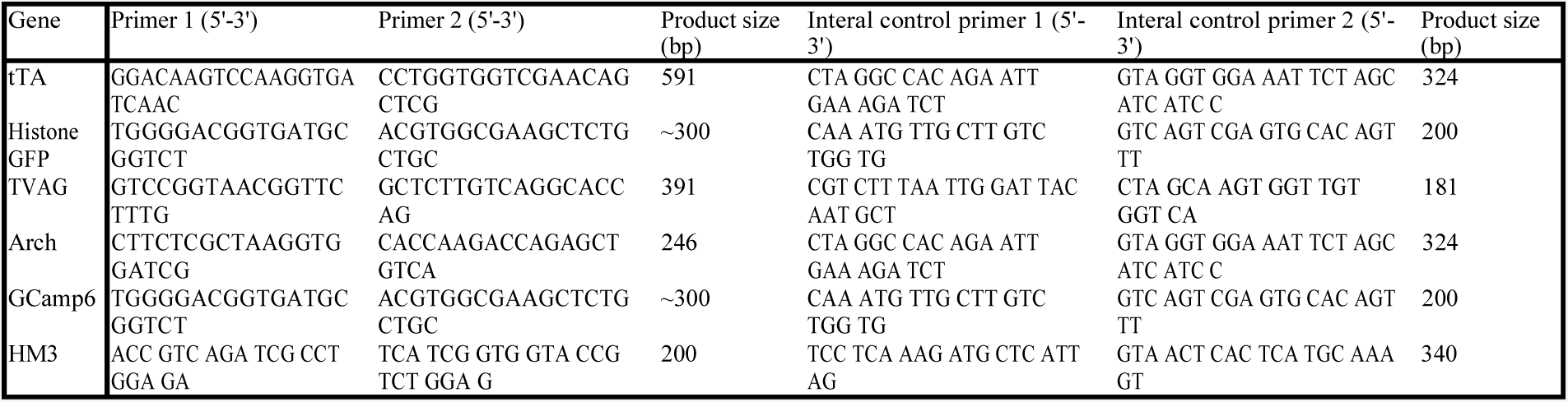
Primers used for genotyping, related to STAR methods. All primers were used in a concentration of 10µM. All genes are genotyped individually, ie. tTA x TVAG crosses were genotyped using two separate reactions, one for tTA and one for TVAG.

